# Structural basis for conserved and distinct antigen recognition by a lineage of malaria-protective antibodies

**DOI:** 10.64898/2026.01.30.702933

**Authors:** Monika Jain, Fabien Cannac, Sashank Agrawal, Wen-Hsin Lee, Johannes R. Loeffler, Monica L. Fernández-Quintero, Gonzalo E. González-Páez, Re’em Moskovitz, Andrew B. Ward, Ian A. Wilson

## Abstract

Monoclonal antibodies (mAbs) targeting the *Plasmodium falciparum* circumsporozoite protein (PfCSP) have demonstrated substantial promise in preventing malaria infection and disease. PfCSP is characterized by a central region composed of repetitive NANP motifs, which serve as major targets for protective antibodies. Several potent mAbs targeting this region exhibit homotypic Fab-Fab interactions, which enhance antigen binding and contribute to their neutralization potency. Among these, mAb 399, encoded by the *IGHV3-49/IGKV2D-29* (V_H_3-49/V_K_2D-29) germline lineages, forms head-to-head inter-Fab contacts mediated primarily by germline-encoded residues. In this study, we determined high-resolution X-ray crystal and cryo-EM structures of two additional Fabs, derived from the same germline lineages, 7160 and 7118, in complex with PfCSP-derived peptides. Both Fabs bound the NANP_6_ repeats with high affinity (K_D_ 5-10 nM). Fab 7160 formed similar inter-Fab homotypic interactions to Fab 399 upon binding to an extended repeat peptide (NANP₆), indicating a conserved mode of recognition. In contrast, Fab 7118, which features a longer CDRH3, does not form homotypic contacts. Instead, Fab 7118 induced a bend in the peptide and adopted a distinct binding mode, which prevents inter-Fab interactions. A cryo-EM structure of Fab 7118 in complex with a recombinant shortened PfCSP construct (rsCSP) further revealed potential disruptions at inter-Fab contact sites. These findings highlight the structural versatility of V_H_3-49/V_K_2D-29-derived antibodies and demonstrate that CDR loop variations within a shared antibody lineage can modulate antibody conformation, homotypic Fab-Fab interactions, and epitope engagement. Our study provides mechanistic insights into the diverse strategies by which CSP-specific antibodies can achieve high-avidity binding and protective immunity.

## Introduction

Malaria remains a global concern and is endemic in many tropical countries, particularly in sub-Saharan Africa and Southeast Asia, despite substantial progress that has been made against malaria control over the past decades (1, 2). Recent reports indicate an increase in malaria cases, driven by factors such as climate change, insecticide-resistant mosquitoes, and drug-resistant parasites (3, 4). In 2023, an estimated 263 million cases and 597,000 deaths from malaria were reported worldwide (5). Malaria is a mosquito-borne disease caused by unicellular parasites of the *Plasmodium* species. *Plasmodium falciparum* is the most lethal of these parasites, with a complex life cycle that involves both sexual and asexual stages occurring in humans and mosquitoes (6). Several vaccines have been developed to target malaria, which are broadly categorized based on the *Plasmodium falciparum* life cycle: pre-erythrocytic vaccines, blood-stage vaccines, and transmission-blocking vaccines (7). The circumsporozoite protein (CSP) of *P. falciparum* is the primary antigen targeted in the development of pre-erythrocytic vaccines. *Plasmodium falciparum* circumsporozoite protein (PfCSP) comprises three main regions: an N-terminal domain involved in hepatocyte attachment, a central immunodominant repeat region (comprising one NPDP, four NVDP, and approximately 35–41 NANP repeats), and a C-terminal thrombospondin repeat (αTSR) domain that mediates sporozoite attachment (8). Among pre-erythrocytic vaccines, RTS,S and R21 are safe and effective in preventing malaria and have been approved by the World Health Organization (WHO) (9). Both vaccines are based on a truncated version of CSP, containing 19 NANP repeats from the central region and the C-terminal αTSR domain. However, their efficacy declines over time (10–13). One proposed explanation for this waning immunity is that the CSP repeat region may not effectively engage B cells, leading to the generation of predominantly short-lived plasmablasts rather than long-lived memory B cells (14, 15).

Over the past few years, several human monoclonal antibodies (mAbs) specific to PfCSP have been isolated and shown to be protective in preclinical models (16). Notably, highly potent mAbs such as CIS43, L9, and MAM01, each targeting different regions of CSP, specifically the junctional, minor repeat, and central repeat regions, respectively, have completed Phase I clinical trials (17–20). Several high-resolution structures of antigen-binding fragments (Fabs) of protective monoclonal antibodies (mAbs) in complex with different regions of PfCSP have been elucidated using X-ray crystallography and cryo-electron microscopy (21–23). Notably, a subset of mAbs encoded by diverse germline genes, including antibodies 850, 1210, 1450, 311, and 399, form Fab-Fab homotypic contacts upon binding to the central NANP repeat region and exhibit potent neutralizing activity (24–27). Moreover, V_H_3-33-derived mAbs adopt a range of extended helical or spiral conformations when complexed with rsCSP, which is a recombinant shortened PfCSP construct containing a reduced number of NANP/NVDP repeats: 19/3 repeats instead of 38/4 for the *P. falciparum* 3D7 strain. The homotypic interactions between adjacent Fabs help stabilize this extended spiral configuration, enhancing binding avidity and potentially contributing to their protective efficacy (28). In contrast, mAb 399, derived from the V_H_3-49/V_K_2D-29 germline lineage, has been reported to bind *Plasmodium falciparum* circumsporozoite protein (PfCSP) via inter-Fab homotypic contacts arranged in a head-to-head configuration (25). Notably, in this case, these contacts are largely mediated by germline-encoded residues, suggesting that antibodies from this lineage may have an innate tendency to form Fab-Fab contacts in the presence of CSP repeat peptides even without somatic hypermutation.

To further investigate this lineage, we examined mAbs 7160 and 7118, isolated from B-cells of protected individuals who participated in a Phase 2a clinical trial of the RTS, S/AS01 malaria vaccine (20, 29). Both these mAbs, 7160 and 7118, derived from V_H_3-49/V_K_2D-29 germline genes, exhibited strong inhibitory activity, reducing liver-stage sporozoite burden by 90–95% in a murine challenge model, and showed high-affinity binding to the central NANP repeat region of PfCSP (20). Sequence alignment with the previously characterized mAb 399 revealed high similarity across all three antibodies. However, mAb 7118 has a slightly different CDRH2 and a distinct CDRH3, which is extended by one additional amino acid residue relative to 399 and 7160 (Fig. 1A).

**Figure 1.**
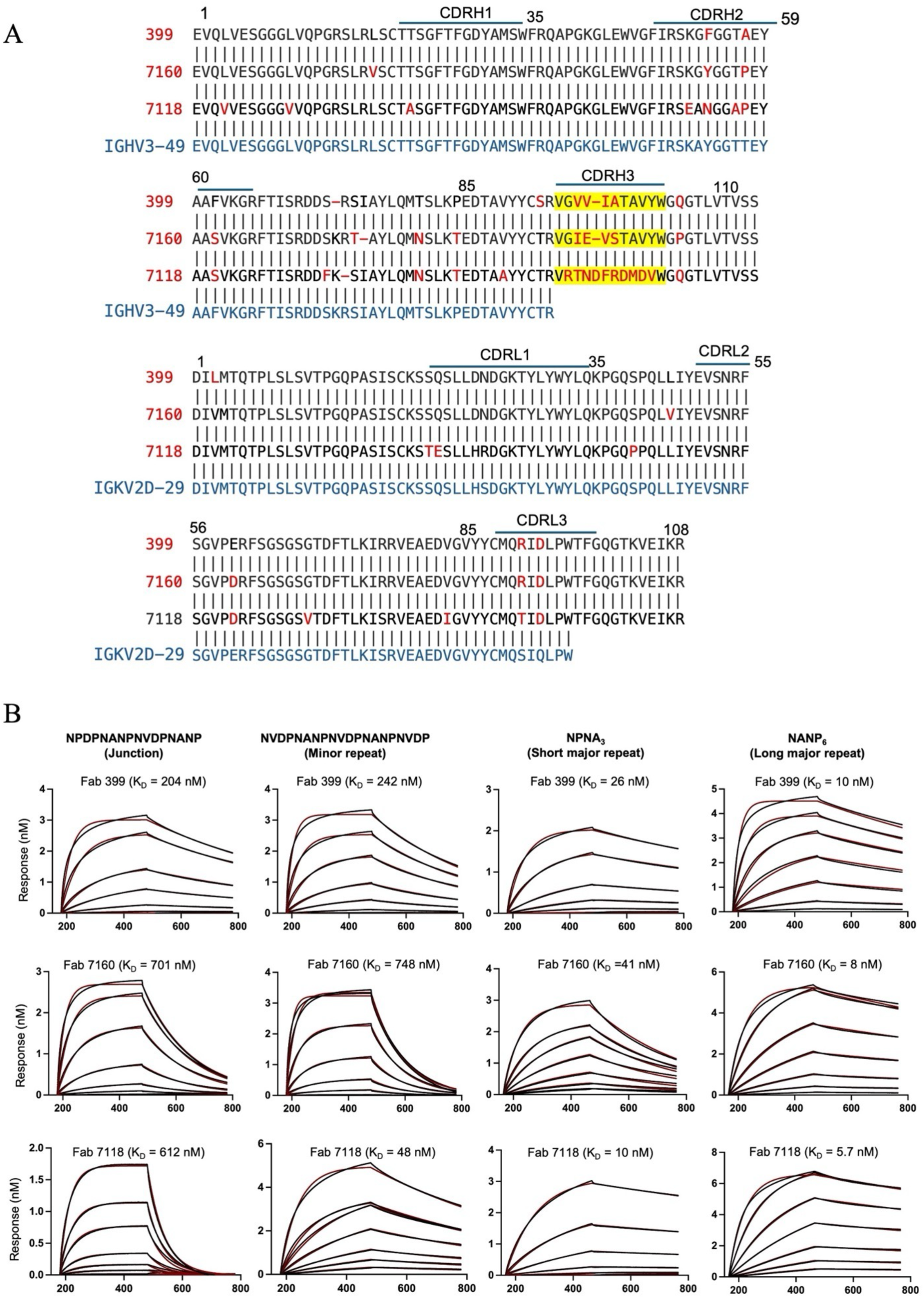
Binding kinetics for Fabs with PfCSP-derived peptides from different regions. **(A)** Sequence alignment of Fabs 7160 and 7118 with 399 reveals a shared V_H_3-49/V_K_2D-29 lineage. Amino acid differences between mAbs 399, 7160, and 7118 are highlighted in red. Sequences of CDRH3 of these mAbs are highlighted in yellow. The alignment between the Fabs heavy/light chain sequence with the germline V_H_3-49/V_K_2D-29 gene sequence is shown (blue) to enable identification of somatically mutated residues. **(B)** Bio-layer interferometry analysis of binding kinetics for V_H_3-49/V_K_2D-29 germline-encoded Fabs 399, 7160, and 7118 with PfCSP-derived peptides of different regions. Binding curves and fitted curves (1:1 global fitting model) are shown in red and black, respectively. Fab concentrations were serially diluted 200, 100, 50, 25, 12.5 nM from top to bottom. Apparent dissociation constants (K_D_) for each Fab-peptide interaction are indicated.

To gain structural and mechanistic insights into this lineage, we determined high-resolution X-ray crystal structures of Fabs 7160 and 7118 in complex with PfCSP-derived peptides, as well as cryo-EM structures of Fabs 7160 and 7118 bound to rsCSP. Structural analysis revealed that Fab 7160 forms inter-Fab homotypic contacts when binding an extended NANP₆ repeat peptide, similar to the interactions observed with Fab 399 (25). In contrast, Fab 7118 did not display such interactions. Instead, its longer CDRH3 loop appears to create steric hindrance and electrostatic repulsion, preventing homotypic Fab-Fab contacts. Additionally, the extended CDRH3 induces a bend in the bound peptide, further differentiating its binding mode from that of 7160 and 399. These findings suggest that even within the same germline lineage, subtle differences in antibody sequence and structure can influence the formation of homotypic Fab-Fab interactions and interaction with the peptide. Overall, our results highlight the structural flexibility of V_H_3-49/V_K_2D-29-derived antibodies and provide new molecular insights into how CSP-specific antibodies achieve high avidity and strong inhibition through different modes of binding interactions.

## Results

### Binding affinity of V_H_3-49/V_K_2D-29-derived potent mAbs 399, 7160, and 7118 to PfCSP-derived epitopes

To investigate the epitope specificity and binding of antibodies derived from V_H_3-49/V_K_2D-29 lineage, we evaluated the binding affinities of Fabs 399, 7160, and 7118 to peptides corresponding to different repeat regions of PfCSP. The antibody sequences of 7160 and 7118 were derived from plasmablasts isolated from protected individuals enrolled in a phase 2a clinical trial of the RTS,S/AS01 vaccine, and both demonstrated protective activity in murine challenge models (20). To evaluate the binding specificities of Fabs 399, 7160, and 7118, we performed Bio-layer Interferometry (BLI) using biotinylated peptides representing the junctional region (NPDPNANPNVDPNANP), minor repeat region (NVDPNANPNVDPNANPNVDP), and major repeats of varying lengths (NPNA₃ and NANP₆). All three Fabs bound the major repeats with high affinity, exhibiting dissociation constants (K_D_) in the range of 10-40 nM for NPNA₃ and 6-10 nM for NANP₆ (Fig. 1B, Table S1). Among them, Fab 7118 exhibited stronger binding to both major repeat peptides, with the lowest K_D_ values. Despite their preference for the major repeats, all Fabs also exhibited cross-reactivity with minor and junctional repeat peptides, but with reduced K_D_’s in the 50-750 nM range. Fab 399 showed approximately 9-fold lower affinity for both junctional and minor repeat peptides compared to NPNA_3_, whereas Fab 7160 exhibited ∼20-fold lower affinity for these regions. Notably, Fab 7118 retained relatively strong binding to the minor repeat (K_D_ = 48 nM) yet displayed much weaker affinity for the junctional region (K_D_ = 612 nM), similar to the other antibodies (Fig. 1B). These results indicate that these V_H_3-49/V_K_2D-29-derived antibodies preferentially bind the major repeat region of PfCSP while exhibiting some cross-reactivity to both junctional and minor repeats.

### Crystal structures of Fab 7160 in complex with the junctional, minor, short major (NPNA_3_), and long major central repeat region (NANP_6_)

To gain mechanistic insights into these V_H_3-49/V_K_2D-29 mAbs, we determined co-crystal structures of Fab 7160 in complex with PfCSP-derived peptides: junctional region, minor repeat, short major repeat (NPNA₃), and long major central repeat (NANP₆) (Fig. 2). These crystals diffracted to resolutions of 2.30 Å, 2.27 Å, 1.90 Å, and 2.35 Å, respectively (Table S2). The interactions between Fab 7160 and the peptides are primarily mediated by the heavy chain complementarity-determining regions CDRH2 and CDRL3, with additional contacts contributed by CDRH3 and, to a lesser extent, through CDRL1 (Fig. 2B and Fig. S1).

**Figure 2.**
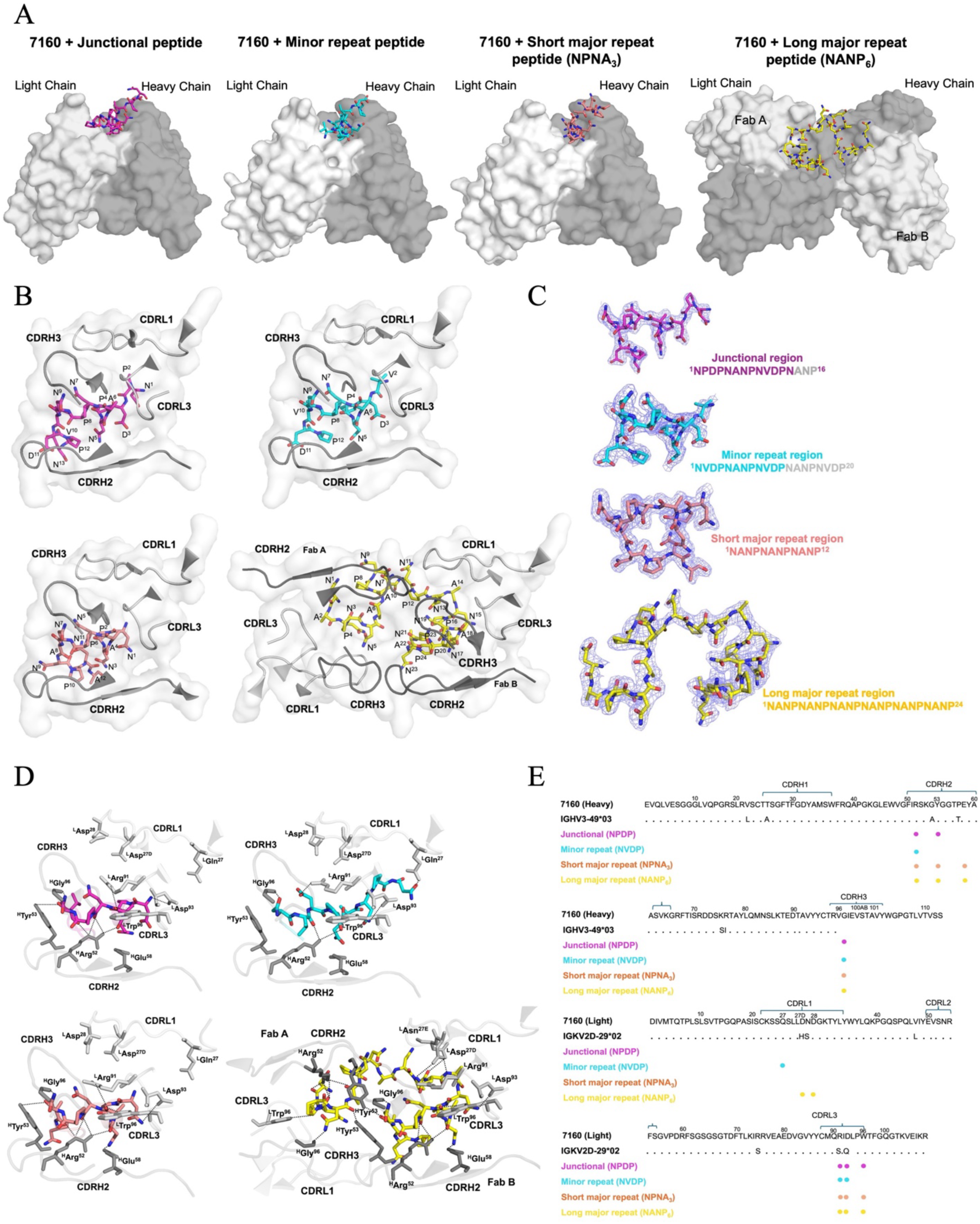
Structural basis of 7160 Fab binding to PfCSP peptides. **(A)** Surface representation of the 7160 Fab in complex with peptides from the junctional (magenta), minor repeat (cyan), short major repeat (salmon), and long major repeat (yellow) regions of PfCSP. The variable domains are colored dark grey (heavy chain) and light grey (light chain). **(B)** Close-up views of 7160-CDRs interacting with each peptide, shown in stick representation and colored as in (A). **(C)** 2Fo–Fc electron density maps (contoured at 2.0σ, shown in grey) for each peptide bound to 7160: junctional (magenta), minor repeat (cyan), short major repeat (salmon), and long major repeat (yellow). Peptides are represented as sticks. **(D)** Hydrogen-bonding interactions between 7160 and each peptide are indicated by black dashed lines. **(E)** Fab 7160 amino acid sequence (Kabat numbering), with CDR residues involved in peptide binding marked by closed colored circles corresponding to each peptide region.

The structure of Fab 7160 in complex with the junctional peptide was crystallized in a P2_1_2_1_2_1_ space group with eight complexes in the asymmetric unit (ASU), whereas the Fab 7160 complex with minor repeat region was also crystallized in the P2_1_2_1_2_1_ space group, but with two complexes per ASU. In the Fab 7160-junctional peptide complex (^1^NPDPNANPNVDPNANP^16^), residues were numbered from 1 to 16. However, electron density was absent for the last three C-terminal residues. In the complex with the minor repeat peptide (^1^NVDPNANPNVDPNANPNVDP^20^), residues were similarly numbered from 1 to 20, but no electron density was observed for residues Asn^13^ to Pro^20^. However, complexes of Fab 7160 with the short major repeat (^1^NPNANPNANPNA^12^) and long major repeat (^1^NANPNANPNANPNANPNANPNANP^24^) exhibited well-defined electron density for all peptide residues (Fig. 2C). Fab 7160 recognizes all peptides in a similar conformation (Fig. 2A and 2B), characterized by alternating type I β-turns and Asn pseudo-3₁₀ turns (Fig. S2). The core of the major repeat region adopts an almost identical conformation to that of the junctional region, with conserved hydrogen bond interactions involving heavy chain ^H^Arg^52^, ^H^Tyr^53^, ^H^Gly^96^, and light chain ^L^Arg^91^, ^L^Asp^93^, and ^L^Trp^96^ (Fig. 2D). The 7160 Fab bound to the minor repeat peptide with fewer hydrogen bonds in CDRH2, but with an additional hydrogen bond involving ^L^Gln^27^ in CDRL1. Consistent with the binding K_D_’s, Fab 7160 forms more extensive hydrogen bonding with the NANP_6_ peptide, and two Fab molecules simultaneously engage a continuous long major repeat epitope, through Fab-Fab homotypic interactions (Fig. 2D and 2E).

### Fab 7160 recognizes short and long NANP repeat epitopes through a conserved binding mode shared with Fab 399

To further analyze the epitope recognition by Fab 7160, we compared the crystal structures of Fab 7160 in complex with short and long peptides derived from the PfCSP central NANP repeat region to the previously reported structures of Fab 399 (25). Fab 7160 with NPNA₃ crystallized in a C2₁ space group with two complexes in the asymmetric unit (ASU), whereas Fab 399–NPNA₃ (PDB ID: 6WFZ) crystallized in a P2₁ space group, also with two complexes per ASU (Fig. 3A). Superimposition of the variable domains of the 7160-NPNA₃ and 399-NPNA₃ complexes revealed a root-mean-square deviation (RMSD) of 0.26 Å across all C-alpha atoms, indicating a nearly identical overall conformation (Fig. 3B). Despite the high structural similarity, Fab 7160 possesses a germline-encoded ^H^Tyr^53^ in CDRH2 that forms additional contacts with the epitope, whereas Fab 399 has ^H^Phe^53^ in the same position. Moreover, the buried surface area (BSA) in the Fab 7160-NPNA₃ complex contributed by the heavy chain CDRH2 and CDRH3 (310 Å² on the Fab and 363 Å² on the peptide) is similar to the Fab 399-NPNA₃ complex (308 Å² on the Fab and 358 Å² on the peptide). Furthermore, in both complexes, the light chain contribution is also substantial and mediated primarily by CDRL3, with the Fab 7160 light chain having a BSA of 215 Å² and 214 Å² on the peptide, comparable to Fab 399 (201 Å² on the Fab and 200 Å² on the peptide) (Fig. 3C). These findings demonstrate that Fab 7160 engages NPNA₃ in a nearly identical manner to Fab 399, suggesting conserved recognition features within this lineage.

**Figure 3.**
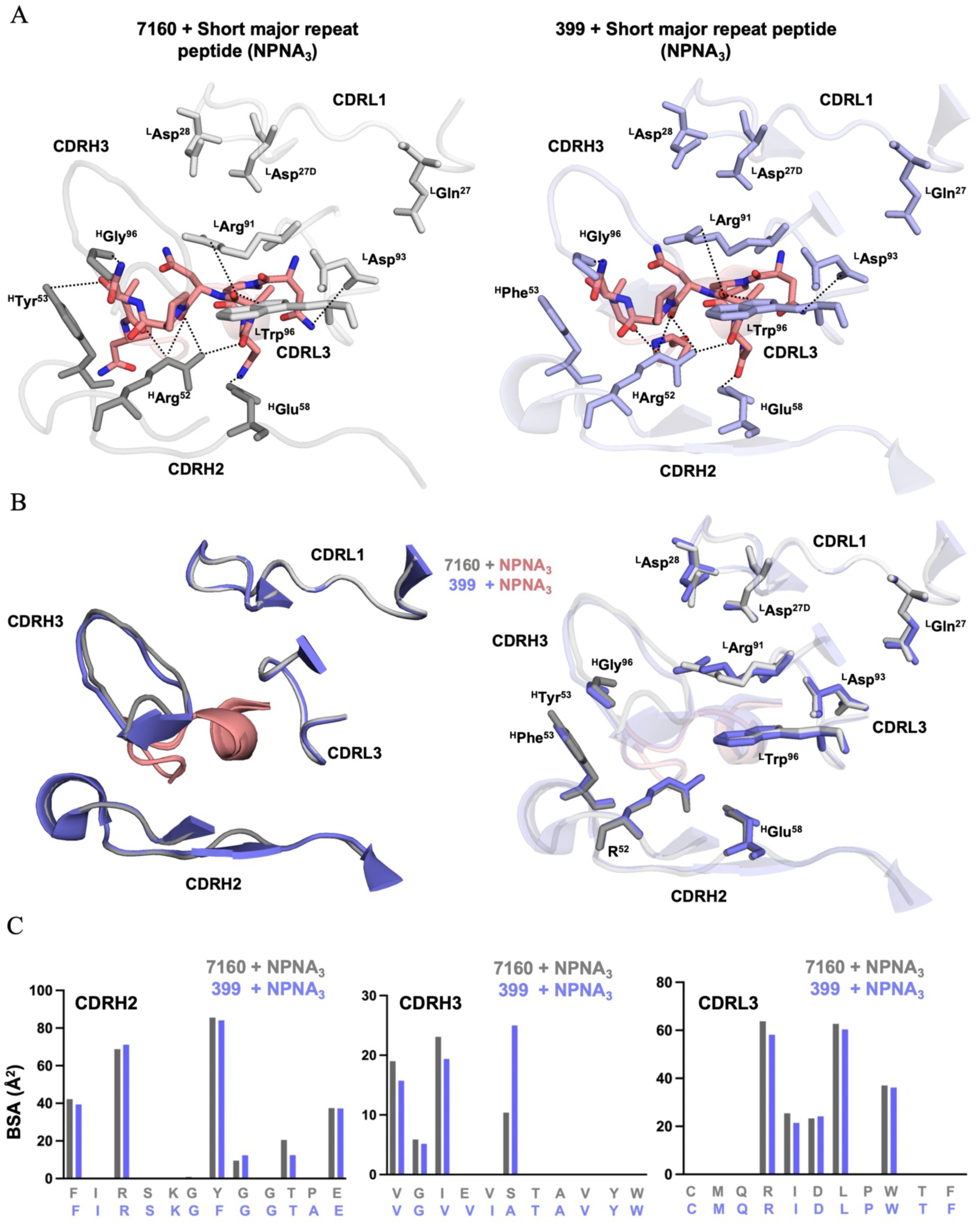
Conserved binding interactions between Fab 7160 and Fab 399. **(A)** Left: Crystal structure of 7160 in complex with NPNA_3_ peptide. Right: Crystal structure of Fab 399 bound to NPNA_3_ (PDB ID: 6WFZ). Only CDRs involved in peptide interactions are shown. Residues forming hydrogen bonds with the peptides are shown as sticks. **(B)** Structural alignment of Fabs 399 (blue) and 7160 (grey) in complex with NPNA_3_ (salmon). **(C)** Bar plot depicting buried surface area contributions from CDRs H2, H3, and L3 in Fab 399 (blue) and 7160 (grey) with bound NPNA_3_ peptide.

Next, we compared the crystal structure of Fab 7160 in complex with the long major repeat region peptide (NANP₆) to that of the previously characterized Fab 399-NANP₆ complex (PDB ID: 6WG1) (25). The 7160-NANP₆ complex crystallized in a P2₁ space group with two complexes in the asymmetric unit (ASU) (Fig. 4A), whereas the 399-NANP₆ complex also crystallized in the P2₁ space group but with only a single complex in the ASU (Fig. 4B). Structural analysis revealed that Fab 7160 exhibited slightly higher binding affinity and a modestly increased number of hydrogen bonds and inter-Fab contacts with NANP₆ compared to Fab 399 (Fig. S3). The Fab-Fab interactions in both Fab 399 and Fab 7160 complexes are symmetric, with CDRH1 and CDRH3 of one Fab engaging CDRH1 of the adjacent Fab (Fig. 4C). Specifically, ^H^Asp^31^ in CDRH1 and ^H^Arg^94^ in CDRH3 form hydrogen bonds with ^H^Tyr^32^ and ^H^Thr^28^, respectively, in the neighboring Fab.

**Figure 4.**
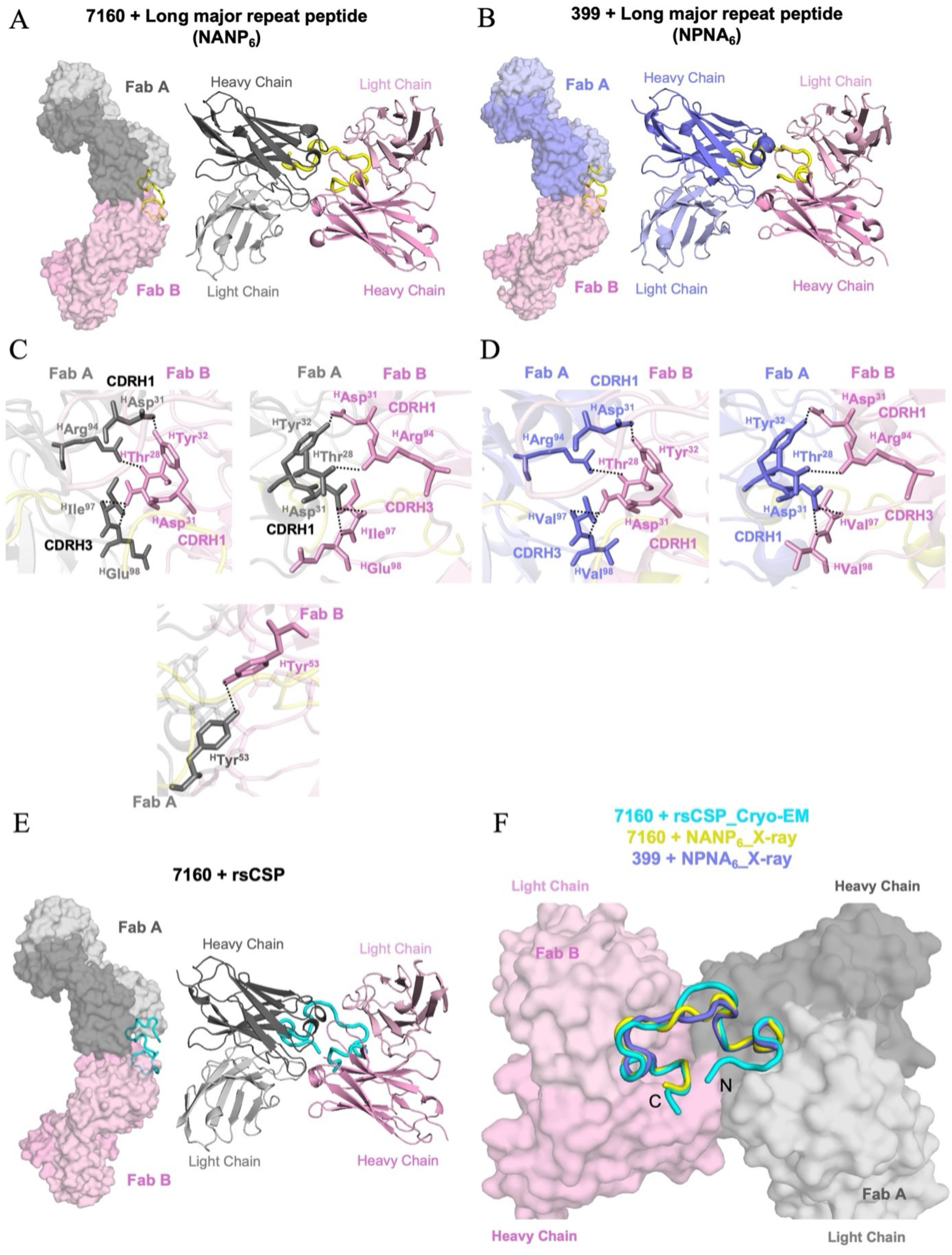
Homotypic Fab-Fab interactions and peptide engagement by Fabs 7160 and 399. **(A)** Homotypic head-to-head interactions between two Fab 7160 molecules are shown in both surface and cartoon representations. Fab A is colored dark grey (heavy chain) and light grey (light chain); Fab B is shown in dark pink (heavy chain) and light pink (light chain). The NANP_6_ peptide is depicted in a yellow cartoon. **(B)** Crystal structure of two Fab 399 molecules bound to NANP_6_ (PDB ID: 6WG1), displayed as surfaces and in cartoon form. Fab A is colored dark blue (heavy chain) and light blue (light chain), and Fab B is colored dark pink (heavy chain) and light pink (light chain). **(C)** Hydrogen-bonding interactions within the homotypic interface formed by two Fab 7160 molecules simultaneously engaging the NANP_6_ peptide. **(D)** Interactions between the two Fab 399 molecules simultaneously engaging an NPNA_6_ peptide. Hydrogen bonds are indicated by black dashed lines. The Fabs are shown as cartoon representations with the side chains of interacting residues represented as sticks. **(E)** Cryo-EM structure of Fab 7160 in complex with rsCSP. Two Fab 7160 molecules bound to rsCSP are shown in surface and cartoon representations, with rsCSP colored cyan. **(F)** Structural overlay comparing the cryo-EM structure of 7160-rsCSP (cyan) with the X-ray structures of Fab 7160 (yellow) and 399 (dark blue) bound to CSP-derived NANP_6_ peptide.

Additionally, ^H^Ile^97^ and ^H^Glu^98^ form a network of mainchain-to-sidechain hydrogen bonds with ^H^Asp^31^ of the adjacent Fab. A similar hydrogen-bonding pattern was observed in the Fab 399-NANP_6_ complex (Fig. 4D). Importantly, ^H^Tyr^53^ in Fab 7160 plays a dual role by mediating key homotypic Fab-Fab contacts and directly participating in CSP binding (Fig. 4C and 2B). Overall, Fab 7160 adopts a mode of recognition nearly identical to that of Fab 399 for both short and long NANP repeat epitopes, characterized by conserved epitope contacts and symmetric, head-to-head Fab–Fab interactions. This binding configuration represents a unique binding strategy among PfCSP-directed antibodies, as the formation of homotypic contacts is symmetric and is largely independent of somatic hypermutation (SHM). This contrasts the spiral arrangement adopted by antibodies derived from the V_H_3-33 germline that are predominantly expressed among PfCSP-binding antibodies, highlighting diverse lineage-specific modes of antigen recognition.

### Cryo-EM of Fab 7160 in complex with recombinant rsCSP

Both the previously reported Fab 399-NANP_6_ and our new Fab 7160-NANP_6_ crystal structures display a head-to-head interface with a relatively short peptide bound in between the two Fab molecules, posing the question of whether a longer antigen, such as CSP, would accommodate such a tight interface. To assess whether the Fab 7160 interface allows a CSP antigen to fully occupy the binding sites on both Fab molecules simultaneously, we solved the cryo-EM structure of Fab 7160 in complex with rsCSP, as previously described (28). Both X-ray and cryo-EM structures of the Fab dimers are nearly identical, with an overall RMSD of 0.47 Å (Fig. S4A). In the cryo-EM structure, however, both epitopes are fully occupied by rsCSP with density for 31 antigen residues observed (vs. 24 for the crystal structures), confirming that the head-to-head interface of the variable domains does not hinder binding of both Fab molecules to the CSP antigen. As expected from the symmetrical Fab-Fab interface, antigen binding is also symmetrical (Fig. 4E), with the N and C-termini of the bound NANP repeats in the rsCSP exiting the complex on the same side. The short linker region between adjacent CSP epitopes is flexible and adopts different conformation in the crystal and cryo-EM structures (Fig. 4F and Fig. S4A), as it enables each epitope to orient correctly for Fab binding on either side. Furthermore, although we could not obtain 3D models for these, we observed 2D classes indicating a higher-order arrangement of the Fabs with oligomerization of the dimers (Fig. S4B). These are clear indications that multiple Fab-7160 complexes can coexist on one CSP antigen, and more investigation will be required to solve their 3D cryo-EM structures, as these higher oligomers suffer from a strong orientation bias.

This cryo-EM work represents the first complex structure of an antibody from this lineage with a longer CSP molecule and confirms that, despite the tight space at the Fab-Fab interface, such antibodies are capable of binding the antigen as it would be presented on the parasite, while retaining their homodimeric form and with multiple antibodies binding the same antigen.

### Crystal structures of Fab 7118 reveal a unique binding mode for the PfCSP repeat region

To investigate the structural basis of Fab 7118 recognition of the PfCSP, we determined X-ray structures of Fab 7118 in complex with peptides derived from the minor repeat region and long major central repeat region of PfCSP. The crystals of both complexes diffracted to high resolutions of 2.09 Å and 1.99 Å, respectively (Table S3). Both complexes crystallized in space group P2_1_2_1_2_1,_ each with one Fab-peptide complex per asymmetric unit (ASU) (Fig. 5A and 5B). The interactions between Fab 7118 and the peptides are mediated by CDRH2, CDRH3, CDRL1, and CDRL3. In the 7118 Fab complex with the minor repeat peptide (^1^NVDPNANPNVDPNANPNVDP^20^), the electron density for Asn^1^ was poorly defined (Fig. 5C). In the complex with long major repeat (^1^NANPNANPNANPNANPNANPNANP^24^), electron density was absent for the first N-terminal and last three C-terminal residues (Fig. 5D). The core of the major repeat region adopts an almost identical conformation to that of the minor repeat region (Fig. 5E and 5F). The conserved hydrogen bond interactions to the peptides involve heavy chain ^H^Arg^52^, ^H^Asn^53^, ^H^Glu^58^, ^H^Arg^96^, ^H^Asn^98^ and ^H^Phe^100^, and light chain ^L^Glu^27^, ^L^Ser^27A^, ^L^Leu^27C^, ^L^Arg^27E^, ^L^Asp^93^, and ^L^Trp^96^ (Fig. 5G). Unlike Fabs 399 and 7160, Fab 7118 does not form inter-Fab homotypic interactions. Instead, 7118 engages in extensive hydrogen bonding with its peptide epitope. In particular, CDRH3 and CDRL1 of 7118 form an additional and distinct set of hydrogen bond contacts compared to those observed in 399 or 7160 (Table S4A and S4B). These additional interactions in Fab 7118 likely compensate for the absence of inter-Fab stabilization, helping to maintain high-affinity binding. Structural superimposition of Fab 7118 and Fab 7160 in complex with NANP_6_ highlights a distinct peptide-binding mode in 7118, where all interactions with the peptide are accommodated within a single Fab (Fig. S5). Notably, the CDRH3 loop in 7118 is longer and contains charged (Arg, Asp) as well as bulky (Phe) amino acid residues (Fig. S5A). These features likely introduce steric hindrance and electrostatic repulsion, thereby preventing head-to-head Fab-Fab contacts observed in 7160 and 399 (Fig. S5B and S5C). This extended CDRH3 induces a bend in the bound peptide, suggesting an alternative mechanism for achieving affinity and stabilizing the Fab-epitope complex in the absence of homotypic interactions. Nevertheless, despite these differences in peptide-binding mode, all three Fabs recognize a conserved and similar binding mode for two NANP repeats.

**Figure 5.**
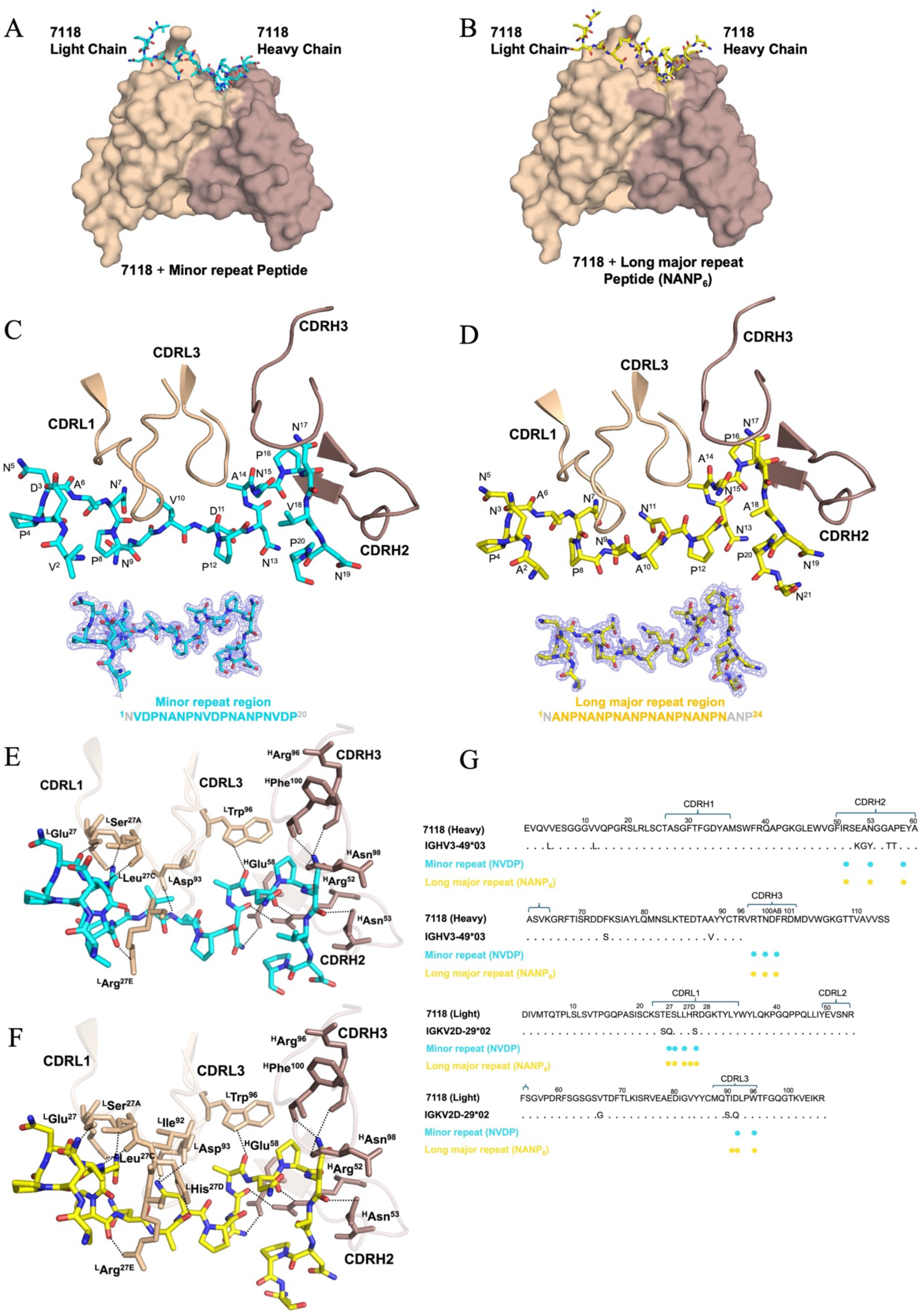
Structural basis of 7118 Fab binding to minor and major repeat peptides. **(A-B)** Surface representation of the 7118 Fab in complex with peptides from the minor repeat (cyan) and long major repeat (yellow) regions of PfCSP. The variable domains are colored brown (heavy chain) and wheat (light chain). **(C-D)** Close-up views of 7118-CDRs interacting with each peptide, shown as stick representation and colored as in panels A and B. 2Fo–Fc electron density maps (contoured at 2.0σ, shown in grey) for each peptide bound to 7118: minor repeat (cyan), and long major repeat (yellow). Peptides are represented as sticks. **(E-F)** Details of hydrogen-bonding interactions between 7118 and each peptide are shown as black dashed lines. **(G)** Amino acid sequence of Fab 7118 (Kabat numbering), with CDR residues involved in peptide binding marked by closed colored circles corresponding to the peptide region.

### Cryo-EM and molecular dynamics simulations of Fab 7118 bound to rsCSP demonstrate a quaternary spiral structure with transient homotypic contacts

To determine how Fab 7118 binds to CSP, we determined a cryo-EM structure of Fab 7118 in complex with rsCSP. The cryo-EM map was resolved to an overall resolution of 3.32 Å (Table S5), allowing visualization of the Fab variable domains bound to repetitive NANP motifs within the central repeat region of rsCSP (Fig. 6A). The structure revealed that four Fab molecules simultaneously bind to the repeating epitopes within rsCSP but without engaging in direct inter-Fab homotypic contacts.

**Figure 6.**
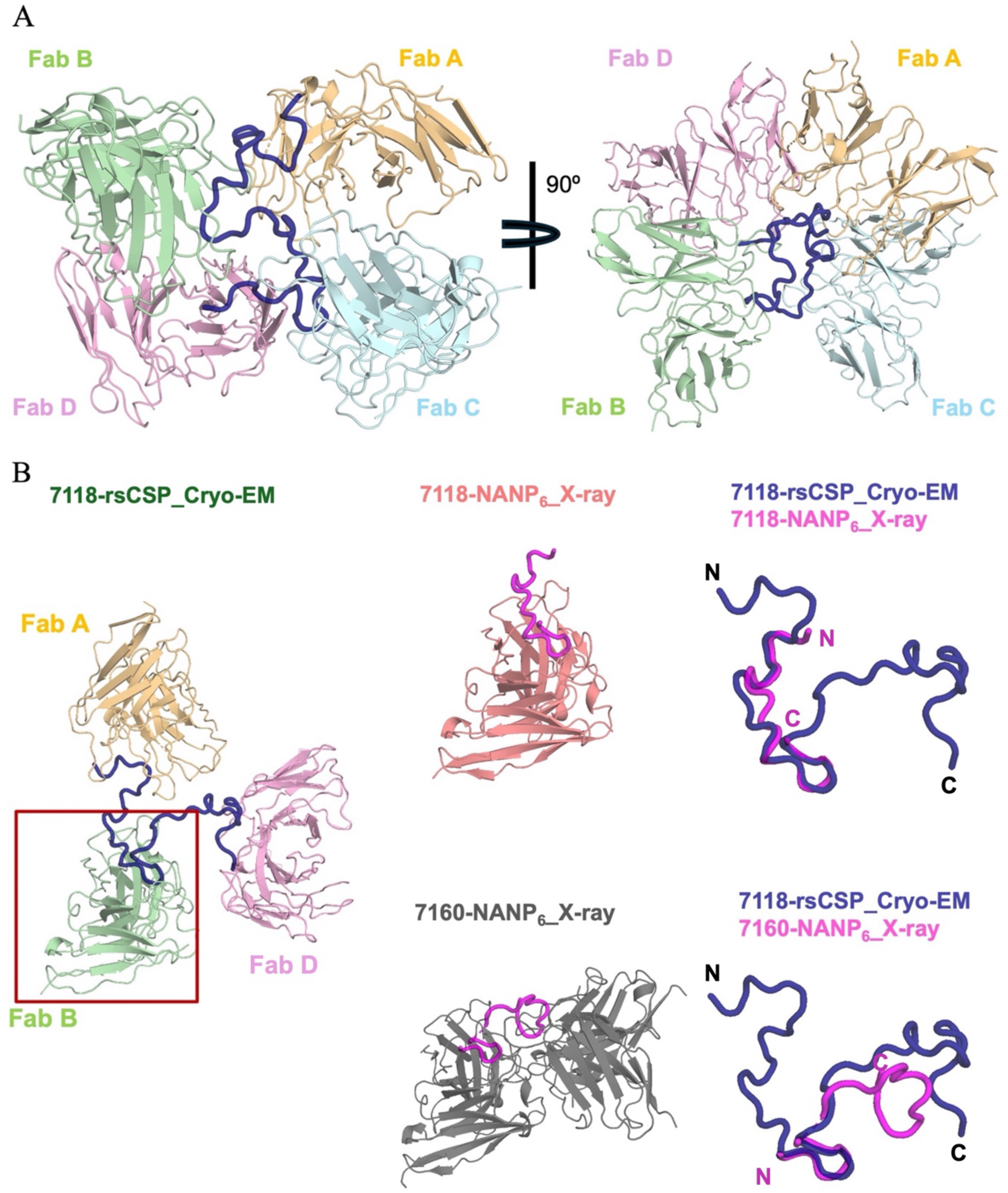
Cryo-EM structure of the 7118 Fab-rsCSP complex. **(A)** Cryo-EM structure of 7118-rsCSP complex at 3.32 Å resolution. A cartoon representation of the atomic model is shown; only the Fab variable regions (Fv) were built into the density. Individual Fabs are colored (Fab A: wheat, Fab B: green, Fab C: cyan, Fab D: pink) and rsCSP is represented as a blue backbone tube. The right panel shows a rotated view. **(B)** Comparison and superposition of Fab variable regions from the 7118-rsCSP cryo-EM structure (green; only Fabs A, B, and D shown) with the 7118-NANP_6_ crystal structure (salmon, upper right) and two Fab molecules from the 7160-NANP_6_ crystal structure (grey, lower right), highlighting differences in peptide binding mode. Models were aligned in PyMOL (53) and displayed in cartoon representation to highlight variations in how the Fabs recognize the NANP repeats.

Structural superposition of the cryo-EM model with the Fab 7118-peptide crystal structure showed minimal conformational differences in the Fab variable domain and bound peptide. This structure contrasts with Fab 7160, which adopts a different peptide-binding orientation in its crystal structure and exhibits inter-Fab contacts (Fig. 6B). When the Fab 7118 cryo-EM structure was superimposed onto the Fab 7160-NANP_6_ complex, we observed steric clashes at potential inter-Fab contact sites, indicating structural incompatibility. These clashes are primarily localized to the longer CDRH3 loop of Fab 7118, particularly at ^H^Asp^99^, which likely introduces steric hindrance that prevents homotypic Fab-Fab interactions and is predicted to interfere with the Fab-Fab packing seen in 7160 and 399 (Fig. S6). Collectively, these findings highlight the unique binding configuration of Fab 7118, where epitope engagement compensates for the lack of inter-Fab interactions, enabling strong and specific recognition of the PfCSP repeat region.

While this binding mode supports the crystal structure data and confirms that Fab 7118 maintains a monovalent mode of engagement with each repeat epitope, we investigated the stability of the quaternary structure observed in the absence of strong homotypic contacts. We performed 3D Variability (3DVA) (30) on the cryo-EM data using particles from the final reconstruction, and the resulting volumes described a homogeneous spiral despite the lack of apparent homotypic contacts, with most of the motion occurring as individual Fab molecules pivot around a constrained CSP spiral (Fig. 7A).

**Figure 7.**
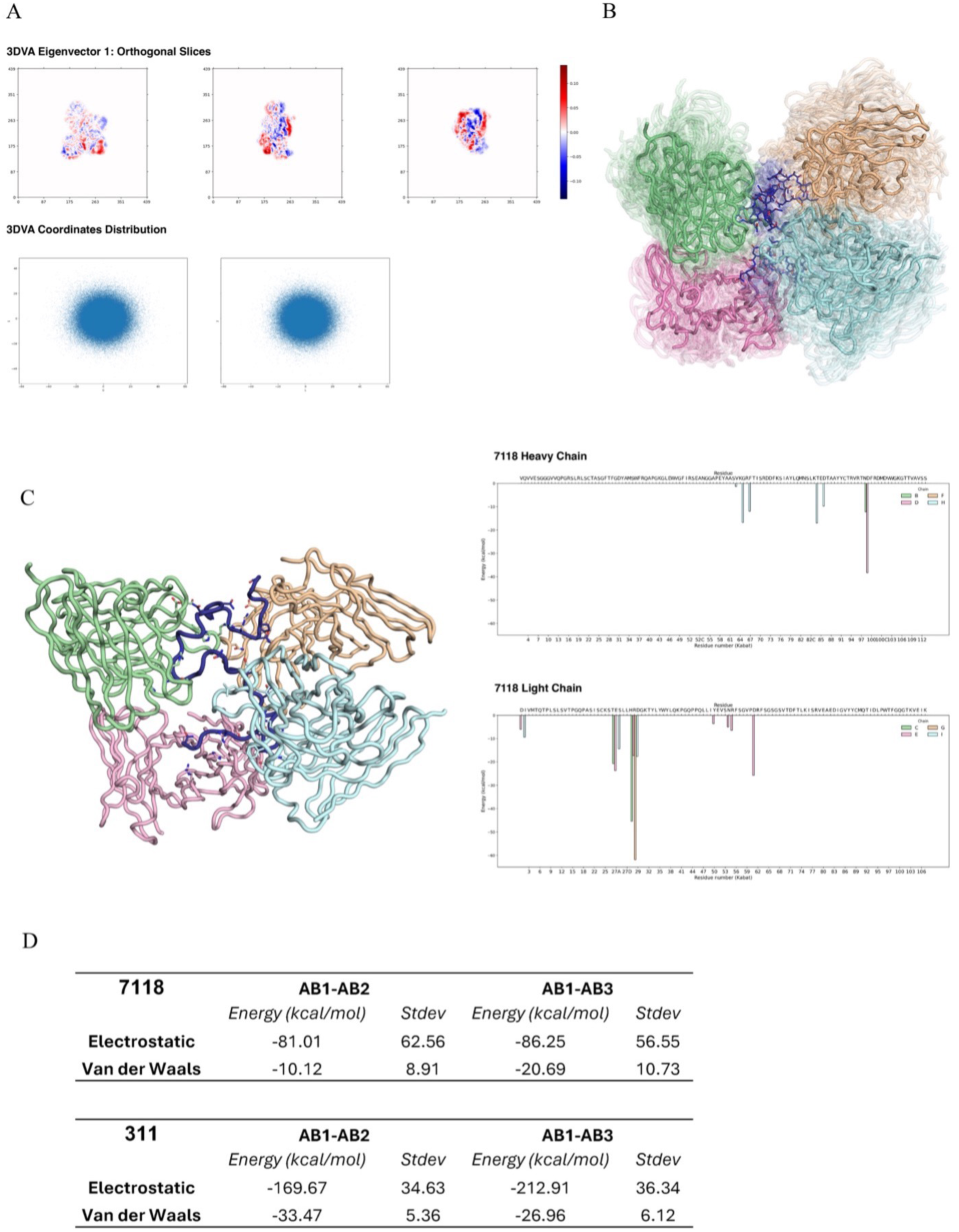
3D Variability Analysis and molecular dynamic simulations of 7118-rsCSP complex. **(A)** Top: Orthogonal slices for one eigenvector from 3D Variability Analysis (3DVA) of 7118-rsCSP complex. Bottom: Coordinates distribution along all three eigenvectors from the 3DVA. The relatively weak Fab movements and uniformity of the coordinates confirm the observed spiral to be a stable complex. **(B)** Overlay of all clusters from the molecular dynamics simulations. All three 1 μs simulations were averaged into one trajectory, itself split into 40 clusters based on atom coordinate standard deviations. The most populated cluster is represented as a solid ribbon, and the other clusters as transparent ribbons. All four 7118 Fabs (1: wheat, 2: green, 3: cyan, 4: pink) are represented as ribbons, rsCSP (blue) as sticks with heteroatoms colored (dark blue: nitrogen, red: oxygen). **(C)** Transient 7118 homotypic contacts observed in MD (color code as above). Left: depiction of every residue detected as forming a temporary homotypic contact, with residues involved in transient contacts shown as sticks. Right: Electrostatic energy of contact for each residue position and each Fab, following the color code above. **(D)** Comparison of Fab-Fab energy between 7118 and 311 cryo-EM models, averaging the calculated energy for each complex over the course of the molecular dynamic simulations.

To further characterize the interaction networks and energetics involved in spiral formation and stabilization, we performed molecular dynamics (MD) simulations on our cryo-EM model. The MD simulations revealed high flexibility of the complex reflected in a high conformational diversity, in line with the motions observed in 3DVA (Fig. 7B). Next, we calculated the respective interaction energies between adjacent Fab molecules over the whole simulation trajectory; for each interacting residue the average energy was calculated and the resulting value plotted (Fig. 7C), revealing transient hydrogen-bond homotypic contacts between Fab molecules (Fig. 7C). To define the strength of these homotypic contacts, we compared the average energy at the transient interface of three consecutive 7118 Fab molecules with three Fab molecules from a classic homotypic spiral like 311 Fab (PDB: 8DYX, Fig. 7D). Calculations indicate that the overall electrostatic energy for 311 Fab-Fab interfaces is more than twice that of 7118 with only half the standard deviation. A similar trend is observed for van der Waals energies, largely driven by hydrophobic residues, none of which are involved in transient interactions in 7118. While the energies of the homotypic interface interactions are relatively weak, the CSP-binding energies range from -120 and -139 kcal/mol for three of the antibodies, confirming the strong epitope/paratope interactions (Table S6). The density at the C-terminal end of the spiral was too weak to confidently build the full peptide in the binding site, reflecting the weaker calculated binding in the simulation for this Fab. Altogether, these data indicate that while 7118 complexed with CSP forms a quaternary structure with a spiral conformation, it does not adopt the homotypic states described previously. Whereas the stability of a classic homotypic spiral, such as with antibody 311, is mediated through strong Fab-Fab electrostatic and van der Waals interactions (26, 28), 7118 does not present such contacts, and as such, only transient interactions occur through a handful of residues on the heavy and light chain in the simulations. The 7118 antibody could therefore possibly represent a maturation intermediate of a CSP-binding antibody on the way to acquiring the somatic hypermutations necessary to form a more permanent homotypic state. Therefore, it will be interesting to identify more antibodies from the V_H_3-49/V_K_2D-29 germline to identify maturation characteristics and somatic hypermutation associated with spiral formation.

### Germline-encoded interactions of aromatic Fabs 7160, 7118, and 399

Structural comparisons of Fab 7160 and Fab 7118 bound to PfCSP-derived peptides revealed conserved aromatic interactions that were also present in the Fab 399-NPNA₃ complex (25). In all three Fab-peptide complexes, a phenylalanine at position 50 in CDRH2, encoded by the *IGHV3-49* germline gene, forms CH/π interactions with the conserved proline residue within the NPNA motif (Fig. 8A). This interaction is further supported by a germline-encoded tryptophan at position 96 in CDRL3 (V_K_2D-29), which forms a hydrogen bond with the backbone carbonyl of the alanine residue preceding the type I β-turn (Fig. 8B). Notably, two NANP repeats in these Fabs bind identically, adopting a similar configuration and exhibiting conserved CH/π interactions with their epitopes.

**Figure 8.**
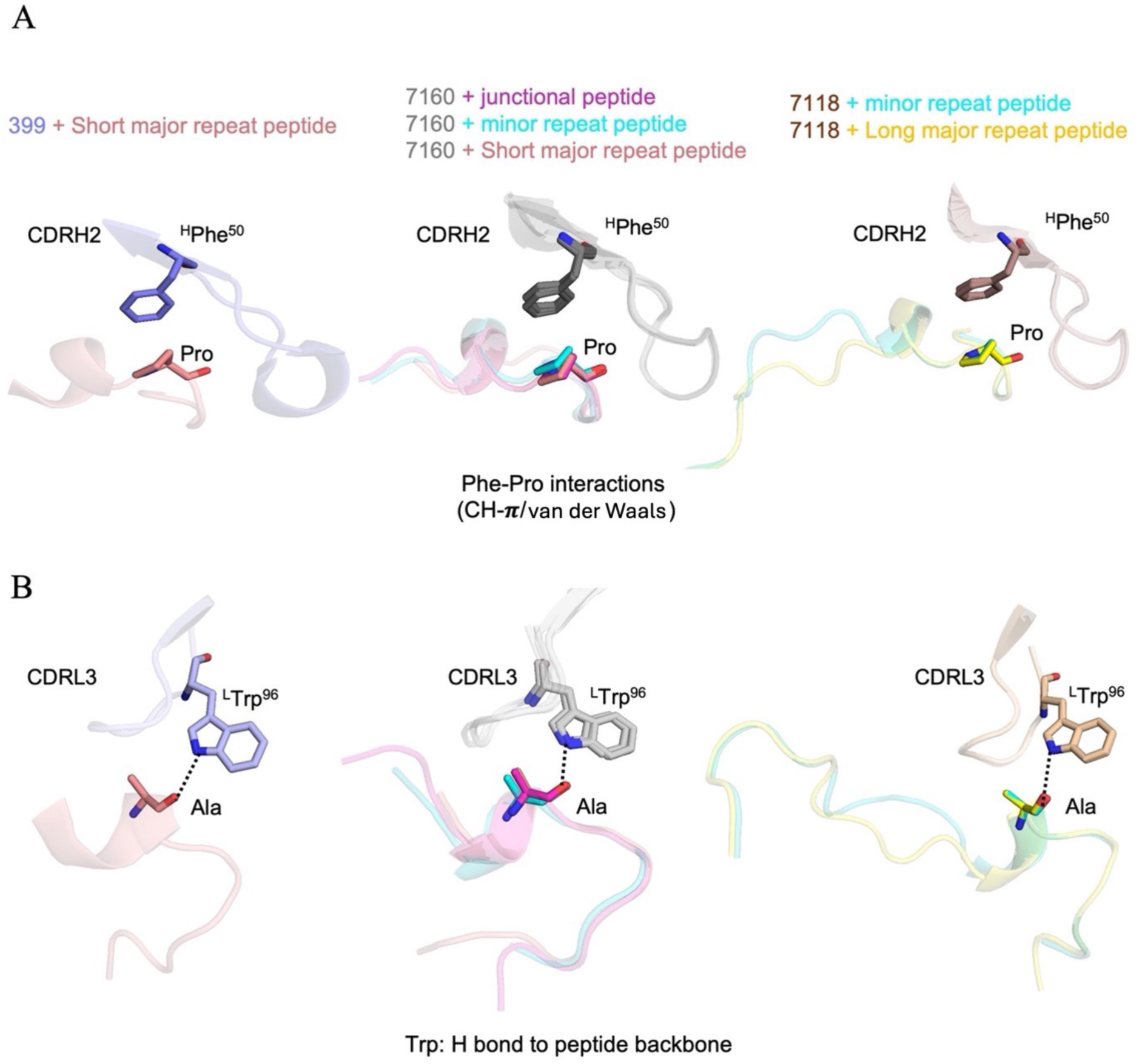
Interactions between Fab aromatic residues and CSP-derived peptides. **(A)** The Fab 399 aromatic residues (blue sticks) interacting with the NPNA_3_ peptide (salmon sticks). **(B)** Superimposed view of 7160 Fab aromatic residues (grey sticks) interacting with peptides from the junctional (magenta), minor repeat (cyan), and short major repeat region (salmon). **(C)** Superimposed view of Fab 7118 aromatic residues (brown sticks) in complex with peptides from minor repeat (cyan), and major repeat region (yellow). Hydrogen bonds are represented as black dashed lines. Residue identities and numbers are labeled with H and L, denoting heavy and light chain, respectively.

These features are consistently observed in the high-resolution crystal structures of 7160 and 7118 bound to peptides from the junctional, minor, and long major repeat regions of PfCSP. Together, these structural findings indicate that these V_H_3-49/V_K_2D-29 Fabs utilize a conserved set of aromatic and hydrogen-bonding interactions to recognize the PfCSP-derived epitopes across different regions.

## Discussion

Our structural characterization of PfCSP-specific mAbs derived from the V_H_3-49/V_K_2D-29 germline lineage provides insights into how antibodies from the same genetic origin can achieve high-affinity binding and potent protection through diverse binding strategies. Fab 7160, like Fab 399, forms inter-Fab homotypic contacts in a symmetric head-to-head configuration upon engaging long NANP repeats. These interactions appear to be germline-encoded and are primarily mediated by heavy chain residues within CDRH1 and CDRH3, with contributions from CDRH2 and CDRL1. In contrast, Fab 7118, despite sharing the same germline origin and exhibiting comparable or even higher affinity for PfCSP repeat peptides, adopts a different binding mode. Structural analysis reveals that Fab 7118 binds in a spiral configuration without permanent inter-Fab contacts, although MD simulation suggests transient contacts between the Fab molecules, which may represent an intermediate state captured in the cryo-EM ensemble. The elongated CDRH3 loop introduces steric hindrance and potential electrostatic repulsion that likely prevents stable inter-Fab contacts, while simultaneously inducing a bend in the bound NANP₆ peptide. These features were consistently observed in both crystal and cryo-EM structures of Fab 7118, confirming that its distinct binding mode is preserved in the context of the recombinant shortened rsCSP antigen. Molecular dynamics simulations to characterize the energy of Fab 7118-rsCSP quaternary structure further revealed transient homotypic contacts that are weaker than those of published homotypic spirals like Fab 311 (26, 28). These results highlight the structural versatility of *IGHV3-49*-derived antibodies, suggesting that not all antibodies from this lineage adopt a similar binding mode, despite their similar antigen specificity. These findings underscore the significance of structural diversity within antibody lineages and highlight the critical role of CDRH3 in governing homotypic interactions. The presence or absence of Fab-Fab contacts and their contribution to overall binding avidity and neutralization may be influenced by subtle sequence differences within CDRs, even among antibodies derived from the same germline genes. Notably, the head-to-head inter-Fab arrangement observed in Fab 7160 and 399 complexes differs from the previously characterized PfCSP-targeting antibodies, derived from the V_H_3-33 germline, which adopt helical or spiral arrangements stabilized by homotypic Fab-Fab contacts (28). Head-to-head inter-Fab interactions have also been reported for V_H_3-33 Fab 1210 and V_H_3-23 Fab 1410 in complex with NANP_5_ (27). However, in these cases, the homotypic interactions are asymmetric and mediated by somatically mutated residues, in contrast to the symmetric, predominantly germline-encoded contacts seen in Fab 7160 and Fab 399. This distinction suggests that different germline-encoded strategies have evolved to engage the repetitive PfCSP epitope, with potential implications for B cell activation, antigen presentation, and the durability of antibody responses, with important implications for vaccine design and efficacy.

From a functional perspective, the ability of all three mAbs-399, 7160, and 7118 to inhibit liver-stage parasite burden by over 90% (20, 25) indicates that both homotypic and non-homotypic modes of binding can provide strong protection. This finding suggests that, while homotypic Fab-Fab interactions may enhance antibody avidity and stability, they are not strictly required for protective efficacy if other factors can compensate. Our data suggest that CSP-targeting antibodies can achieve high avidity and protective efficacy through multiple strategies, either via multivalent epitope engagement stabilized by Fab-Fab interactions or through extensive Fab-epitope hydrogen bonding in the absence of homotypic contacts. In conclusion, our study explains how antibodies of the V_H_3-49/V_K_2D-29 lineage can achieve high-avidity binding to the PfCSP central repeat region through different binding mechanisms. These findings expand our understanding of antibody diversity in response to repetitive antigens and provide valuable guidance for the design of next-generation malaria vaccines targeting PfCSP and similar pathogens.

## Methods and Materials

### Protein expression and purification

The variable region sequences of the heavy and light chains for antibodies 7160 and 7118 were obtained from a published study by Atreca (20). Fab genes were codon-optimized for mammalian expression and cloned into the PHCMV3 expression vector (GenScript). Transient transfection was performed in ExpiCHO cells to express the Fabs. Purification was carried out using a HiTrap Protein G HP column (GE Healthcare), followed by size-exclusion chromatography on a Superdex 200 16/90 column (GE Healthcare) in 1× Tris-buffered saline (TBS; 50 mM Tris, pH 7.8, 140 mM NaCl).

### Biolayer Interferometry (BLI) assay for Fab binding kinetics

The binding kinetics of Fabs 399, 7160, and 7118 to various epitopes of PfCSP were evaluated using biolayer interferometry (BLI) on an Octet Red system (Pall ForteBio). Biotinylated peptides were synthesized by Innopep Inc. and diluted to 10 μg/mL in kinetics buffer (1× PBS supplemented with 0.01% BSA and 0.002% Tween 20). Peptides were immobilized on streptavidin-coated biosensors by incubating for 300 s, followed by a 40 s baseline equilibration in kinetics buffer. For the association, Fabs were serially diluted (200, 100, 50, 25, and 12.5 nM) and allowed to bind to the immobilized peptides for 300 s, followed by a 300 s dissociation step in kinetics buffer. Background subtraction was performed using reference sensors in buffer alone to assess non-specific binding. Kinetic parameters were derived using the Octet data analysis software (version 12.2), with data fitted to a 1:1 binding model.

### Fab-peptide complex formation

All CSP-derived peptides used in this study were N- and C-terminally protected with acetyl and amide groups, respectively. The sequences were as follows: Ac-NPDPNANPNVDPNANP-NH₂ (junctional region peptide), Ac-NVDPNANPNVDPNANPNVDP-NH₂ (minor repeat region peptide), Ac-NPNANPNANPNA-NH₂ (short major repeat region peptide), and Ac-NANPNANPNANPNANPNANPNANP-NH₂ (long major repeat region peptide). All peptides were synthesized by Innopep Inc. at >98% purity. Fab 7160 was concentrated to 10 mg/mL in 1X TBS (Tris-buffered saline) using 10K Amicon® ultra centrifugal filters and incubated with the junctional, minor repeat, and major repeat peptides at a 1:5 Fab-to-peptide molar ratio. Similarly, Fab 7118 was concentrated to 10 mg/mL in 1X TBS and incubated with the minor repeat and long major repeat (NANP₆) peptides at a 1:5 molar ratio. All Fab-peptide mixtures were stored at 4 °C overnight.

### X-ray crystallization

All crystallization screening was performed using our high-throughput robotic CrystalMation system (Rigaku) at The Scripps Research Institute (TSRI). Crystals were obtained using the sitting-drop vapor diffusion method at 293 K, with reservoir solutions of 35 µL and drops consisting of 0.1 µL of protein mixed with 0.1 µL of precipitant solution.

Crystals of Fab 7160 in complex with the junctional region peptide were grown in 0.2 M sodium dihydrogen phosphate (pH 4.5) and 20% (w/v) polyethylene glycol (PEG) 3350. The Fab 7160-minor repeat peptide complex crystallized in 0.1 M HEPES (pH 7.0), 1 M lithium chloride, and 20% (w/v) PEG 6000. Fab 7160 co-crystallized with the NPNA₃ peptide in 0.2 M sodium acetate (pH 7.0) and 20% (w/v) PEG 3350. Crystals of the Fab7160-NANP₆ complex were grown in 0.2 M magnesium nitrate (pH 5.8) and 20% (w/v) PEG 3350. All crystals in this group were cryoprotected using 10% ethylene glycol before data collection.

For Fab 7118 complexes, co-crystals with the minor repeat peptide were obtained in 0.1 M HEPES (pH 7.5), 30% (v/v) 1,2-propanediol, and 20% (v/v) PEG 400. Crystals of the Fab 7118-NANP₆ complex were grown in 0.1 M sodium cacodylate (pH 6.5), 0.16 M calcium acetate, 14.4% (w/v) PEG 8000, and 20% (v/v) glycerol. Crystals for both Fab 7160 and Fab 7118 complexes typically appeared within 14-21 days at 293 K.

### X-ray data collection and structure determination

X-ray diffraction data for Fab 7160 in complex with the junctional and minor repeat peptides were collected at the Advanced Light Source (ALS) beamline BL 5.0.2. Data for Fab 7160 in complex with the short (NPNA₃) and long major (NANP₆) repeat peptides were collected at beamline 17-ID-1 (AMX) at the National Synchrotron Light Source II (NSLS-II). Diffraction data for Fab 7118 in complex with the minor repeat and long major repeat peptides were collected at Stanford Synchrotron Radiation Lightsource (SSRL) beamline BL 12-1.

All datasets were processed and scaled using the HKL-2000 package (31). The structures of Fabs 7160 and 7118 were determined by molecular replacement using PHASER (32) with homology models generated by AlphaFold2 (33). Model building was performed in Coot, and structural refinement was carried out using phenix.refine (34, 35). Peptides were manually fitted into the Fo–Fc electron density maps, followed by multiple cycles of refinement in phenix.refine (34). Fab residue numbering follows the Kabat nomenclature. Structure validation was performed using MolProbity (36). Buried surface areas were calculated with MS (37), and hydrogen bonds were analyzed using HBPLUS (38).

### Cryo-EM sample preparation and data collection

To prepare complexes for cryo-EM, 166 μg of Fab 7118 or Fab 7160 was mixed with 10 μg of rsCSP and incubated overnight at 4°C, followed by purification of the complex by size-exclusion chromatography on a Superdex 200 10/300 column (GE Healthcare), concentrated to around 1 mg/mL, and applied on a glow-discharged Quantifoil 1.2/1.3 300-mesh copper grid (Quantifoil). Grids were blotted for 5 sec after a 10 sec wait time in a Vitrobot Mark IV (ThermoFisher) at 4°C and 100% humidity before being plunged in liquid ethane.

Movies were collected on a Falcon 4i detector (ThermoFisher) mounted to a Glacios 2 microscope (ThermoFisher) operating at 200 keV, with nominal magnification of 190,000X and pixel size of 0.718 Å. Data collection was performed through the EPU interface (ThermoFisher), at an expected dose rate of 60 e^-^/Å^2^ and defocus range between -0.6 μm and -1.6 μm, for a total of 2065 movies (Fab 7118) or 2541 (Fab 7160).

### Cryo-EM data processing and model building

CryoSPARC (39) and CryoSPARC Live were used for data processing. Patch Motion Correction and CTF estimation were performed by cryoSPARC Live and movies with a CTF fit higher than 10 Å were rejected from further processing. An initial batch of particles was picked with Blob Picker (minimal diameter 140 Å), resulting after a first round of 2D classification in a set of templates used for Template Picking on all micrographs. After 2D classification and ab-initio model building, a final set of 250,774 particles (Fab 7118) or 116,764 particles (Fab 7160) was selected for a first Non-Uniform Refinement (NU-R), after which Global CTF Refinement was applied before a final NU-R to obtain the final map. Model building was performed in Coot (35), refined using Phenix (34), and validated with MolProbity (36).

### Molecular dynamics simulation

We used the cryo-EM structures of the antibodies 311 (PDB: 8DYX) and 7118 (this paper) in complex with rsCSP as starting structures for molecular dynamics simulations (MD). Starting structures for MD simulations were prepared in Molecular Operating Environment (Chemical Computing Group, version 2024.06 using the Protonate3D tool (40). To neutralize the charges, the uniform background charge was applied, which is required to compute long-range electrostatic interactions. (41, 42). Using the tleap tool of the AmberTools24 package, the structures were soaked in cubic water boxes of TIP3P water molecules with a minimum wall distance of 12 Å to the protein (43, 44). For all simulations, parameters of the AMBER force field 14SB were used (45). We then performed 3 repetitions of each 1 µs of classical molecular dynamics simulations for each complex using Amber24 (43, 44). MD simulations were performed in an NpT ensemble using pmemd.cuda (46). Bonds involving hydrogen atoms were restrained by applying the SHAKE algorithm, allowing a time step of 2 fs (47). The Langevin thermostat was used to maintain the temperature during simulations at 300 K with a collision frequency of 2 ps^−1^ and a Monte Carlo barostat with one volume change attempt per 100 steps (48–50).

The interaction energies were calculated with CPPTRAJ using the interaction energy (LIE) tool (51). The electrostatic interaction energies were calculated for all frames of each simulation and provided the simulation-averages of these interactions. To calculate the interactions and interaction frequencies of the binding interface, we used the GetContacts tool (52). Cluster analysis was performed in CPPTRAJ, aligning and clustering on all Cα atoms using a distance cut-off criterion of 4 Å. PyMOL was used for visualization of the clusters and the interaction networks (The PyMOL Molecular Graphics System, Version 3.0 Schrödinger, LLC.).

## Data availability

The authors declare that all the data supporting the findings of this study are available within the manuscript and supplementary information files. The structure factors and coordinates of Fab 7160 bound to junctional, minor, short major repeat, and long major repeat epitopes have been deposited in the Protein Data Bank (PDB) under accession codes 9ZM7, 9ZM8, 9ZM9, 9ZMA, respectively. Similarly, the structure factors and coordinates of Fab 7118 bound to the minor repeat and long major repeat epitopes have been deposited under PDB accession codes 9ZMB and 9ZMC, respectively. Cryo-EM structures and corresponding density maps generated in this study for 7160-rsCSP have been deposited in the Protein Data Bank (PDB) and Electron Microscopy Data Bank (EMDB), respectively, under accession codes 9ZFY and EMDB-74165. Similarly, Cryo-EM structures and corresponding density maps generated in this study for 7118-rsCSP have been deposited in the PDB and EMDB, respectively, under accession codes 9ZFZ and EMDB-74167.

## Acknowledgements

We thank H. Tien for assistance with the automated robotic crystal screening at The Scripps Research Institute. X-ray diffraction datasets were collected at the National Synchrotron Light Source II (NSLS II) beamline 17-ID-1, Advanced Light Source (ALS) beamline BL 5.0.2., and Stanford Synchrotron Radiation Lightsource (SSRL) beamline BL 12-1. The authors thank the administration staff of the cryo-EM facility at The Scripps Research Institute. This work was supported and funded by Gates Foundation grants (INV-004923 and INV-056202) under a collaborative agreement with The Scripps Research Institute. The conclusions and opinions expressed in this work are those of the author(s) alone and shall not be attributed to the Foundation. Under the grant conditions of the Foundation, a Creative Commons Attribution 4.0 License has already been assigned to the Author Accepted Manuscript version that might arise from this submission.

## Author contributions

**Conceptualization**: Monika Jain, Ian A. Wilson

**Data Curation**: Monika Jain, Fabien Cannac

**Formal analysis**: Monika Jain, Fabien Cannac, Sashank Agrawal, Johannes R. Loeffler, Monica L. Fernández-Quintero, Re’em Moskovitz

**Funding acquisition**: Ian A. Wilson, Andrew B. Ward

**Investigation**: Monika Jain, Fabien Cannac, Sashank Agrawal

**Methodology**: Monika Jain, Fabien Cannac, Sashank Agrawal, Wen-Hsin Lee, Johannes R. Loeffler, Monica L. Fernández-Quintero

**Project Administration**: Monika Jain, Ian A. Wilson

**Resources**: Monika Jain, Fabien Cannac, Wen-Hsin Lee, Gonzalo E. González-Páez

**Supervision**: Ian A. Wilson, Andrew B. Ward

**Validation**: Monika Jain, Fabien Cannac

**Visualization**: Monika Jain, Fabien Cannac, Ian A. Wilson

**Writing-original draft**: Monika Jain, Ian A. Wilson

**Writing**-**review & editing**: All authors

## References

1. H. J. Oladipo et al., Increasing challenges of malaria control in sub-Saharan Africa: Priorities for public health research and policymakers. Ann Med Surg (Lond) 81, 104366 (2022).

2. Q. Li et al., Malaria: past, present, and future. Signal Transduct Target Ther 10, 188 (2025).

3. C. Caminade et al., Climate change and malaria control: a call to urgent action from Africa’s frontlines. Malar J 24, 179 (2025).

4. P. F. Suh et al., Impact of insecticide resistance on malaria vector competence: a literature review. Malar J 22, 19 (2023).

5. P. Venkatesan, WHO world malaria report 2024. Lancet Microbe 6, e101073 (2025).

6. S. Sato, *Plasmodium*-a brief introduction to the parasites causing human malaria and their basic biology. J Physiol Anthropol 40, 1 (2021).

7. Y. A. Tajudeen et al., A landscape review of malaria vaccine candidates in the pipeline. Trop Dis Travel Med Vaccines 10, 19 (2024).

8. L. T. Wang et al., A potent anti-malarial human monoclonal antibody targets circumsporozoite protein minor repeats and neutralizes sporozoites in the liver. Immunity 53, 733–744 e738 (2020).

9. E. A. Hammershaimb, A. A. Berry, Pre-erythrocytic malaria vaccines: RTS,S, R21, and beyond. Expert Rev Vaccines 23, 49–52 (2024).

10. P. E. Duffy, J. P. Gorres, S. A. Healy, M. Fried, Malaria vaccines: a new era of prevention and control. Nat Rev Microbiol 22, 756–772 (2024).

11. F. Zavala, RTS,S: the first malaria vaccine. J Clin Invest 132, e156588 (2022).

12. P. L. Alonso et al., Efficacy of the RTS,S/AS02A vaccine against *Plasmodium falciparum* infection and disease in young African children: randomised controlled trial. Lancet 364, 1411–1420 (2004).

13. M. S. Datoo et al., Safety and efficacy of malaria vaccine candidate R21/Matrix-M in African children: a multicentre, double-blind, randomised, phase 3 trial. Lancet 403, 533–544 (2024).

14. I. A. Cockburn, R. A. Seder, Malaria prevention: from immunological concepts to effective vaccines and protective antibodies. Nat Immunol 19, 1199–1211 (2018).

15. G. E. Weiss et al., The *Plasmodium falciparum-*specific human memory B cell compartment expands gradually with repeated malaria infections. PLoS Pathog 6, e1000912 (2010).

16. K. Miura, Y. Flores-Garcia, C. A. Long, F. Zavala, Vaccines and monoclonal antibodies: new tools for malaria control. Clin Microbiol Rev 37, e0007123 (2024).

17. Y. Flores-Garcia et al., The *P. falciparum* CSP repeat region contains three distinct epitopes required for protection by antibodies in vivo. PLoS Pathog 17, e1010042 (2021).

18. M. R. Gaudinski et al., A monoclonal antibody for malaria prevention. N Engl J Med 385, 803–814 (2021).

19. N. Nekkab, M. A. Penny, Accelerated development of malaria monoclonal antibodies. Cell Rep Med 3, 100786 (2022).

20. K. L. Williams et al., A candidate antibody drug for prevention of malaria. Nat Med 30, 117–129 (2024).

21. E. Thai et al., Molecular determinants of cross-reactivity and potency by VH3-33 antibodies against the *Plasmodium falciparum* circumsporozoite protein. Cell Rep 42, 113330 (2023).

22. T. Pholcharee et al., Diverse antibody responses to conserved structural motifs in *Plasmodium falciparum* Circumsporozoite Protein. J Mol Biol 432, 1048–1063 (2020).

23. N. K. Kisalu et al., A human monoclonal antibody prevents malaria infection by targeting a new site of vulnerability on the parasite. Nat Med 25, 188–189 (2019).

24. I. Kucharska et al., High-density binding to *Plasmodium falciparum* circumsporozoite protein repeats by inhibitory antibody elicited in mouse with human immunoglobulin repertoire. PLoS Pathog 18, e1010999 (2022).

25. T. Pholcharee et al., Structural and biophysical correlation of anti-NANP antibodies with in vivo protection against *P. falciparum*. Nat Commun 12, 1063 (2021).

26. D. Oyen et al., Cryo-EM structure of *P. falciparum* circumsporozoite protein with a vaccine-elicited antibody is stabilized by somatically mutated inter-Fab contacts. Sci Adv 4, eaau8529 (2018).

27. K. Imkeller et al., Antihomotypic affinity maturation improves human B cell responses against a repetitive epitope. Science 360, 1358–1362 (2018).

28. G. M. Martin et al., Affinity-matured homotypic interactions induce spectrum of PfCSP structures that influence protection from malaria infection. Nat Commun 14, 4546 (2023).

29. J. A. Regules et al., Fractional third and fourth dose of RTS,S/AS01 malaria candidate vaccine: A phase 2a controlled human malaria parasite infection and immunogenicity study. J Infect Dis 214, 762–771 (2016).

30. A. Punjani, D. J. Fleet, 3D variability analysis: Resolving continuous flexibility and discrete heterogeneity from single particle cryo-EM. J Struct Biol 213, 107702 (2021).

31. Z. Otwinowski, W. Minor, Processing of X-ray diffraction data collected in oscillation mode. Methods Enzymol 276, 307–326 (1997).

32. A. J. McCoy et al., Phaser crystallographic software. J Appl Crystallogr 40, 658–674 (2007).

33. J. Jumper et al., Highly accurate protein structure prediction with AlphaFold. Nature 596, 583–589 (2021).

34. P. D. Adams et al., PHENIX: a comprehensive Python-based system for macromolecular structure solution. Acta Crystallogr D Biol Crystallogr 66, 213–221 (2010).

35. P. Emsley, B. Lohkamp, W. G. Scott, K. Cowtan, Features and development of Coot. Acta Crystallogr D Biol Crystallogr 66, 486–501 (2010).

36. V. B. Chen et al., MolProbity: all-atom structure validation for macromolecular crystallography. Acta Crystallogr D Biol Crystallogr 66, 12–21 (2010).

37. S. Yang et al., Ogre: A Python package for molecular crystal surface generation with applications to surface energy and crystal habit prediction. J Chem Phys 152, 244122 (2020).

38. I. K. McDonald, J. M. Thornton, Satisfying hydrogen bonding potential in proteins. J Mol Biol 238, 777–793 (1994).

39. A. Punjani, J. L. Rubinstein, D. J. Fleet, M. A. Brubaker, cryoSPARC: algorithms for rapid unsupervised cryo-EM structure determination. Nature Methods 14, 290–296 (2017).

40. P. Labute, Protonate3D: Assignment of ionization states and hydrogen coordinates to macromolecular structures. Proteins: Structure, Function, and Bioinformatics 75, 187–205 (2009).

41. J. S. Hub, B. L. de Groot, H. Grubmüller, G. Groenhof, Quantifying artifacts in Ewald simulations of inhomogeneous systems with a net charge. J Chem Theory Comput 10, 381–390 (2014).

42. T. Darden, D. York, L. Pedersen, Particle mesh Ewald: An N⋅log(N) method for Ewald sums in large systems. J Chem Phys 98, 10089–10092 (1993).

43. D. A. Case et al., AmberTools. J Chem Inf Model 63, 6183–6191 (2023).

44. D. A. Case et al., Recent developments in Amber biomolecular simulations. J Chem Inf Model 65, 7835–7843 (2025).

45. J. A. Maier et al., ff14SB: Improving the accuracy of protein side chain and backbone parameters from ff99SB. J Chem Theory Comput 11, 3696–3713 (2015).

46. T.-S. Lee et al., GPU-accelerated molecular dynamics and free Energy methods in Amber18: Performance enhancements and new features. J Chem Inf Model 58, 2043–2050 (2018).

47. S. Miyamoto, P. A. Kollman, Settle: An analytical version of the SHAKE and RATTLE algorithm for rigid water models. J Comput Chem 13, 952–962 (1992).

48. S. A. Adelman, J. D. Doll, Generalized Langevin equation approach for atom/solid-surface scattering: General formulation for classical scattering off harmonic solids. J Chem Phys 64, 2375–2388 (1976).

49. J. D. Doll, L. E. Myers, S. A. Adelman, Generalized Langevin equation approach for atom/solid-surface scattering: Inelastic studies. J Chem Phys 63, 4908–4914 (1975).

50. J. Åqvist, P. Wennerström, M. Nervall, S. Bjelic, B. O. Brandsdal, Molecular dynamics simulations of water and biomolecules with a Monte Carlo constant pressure algorithm. Chem Phys Lett 384, 288–294 (2004).

51. D. R. Roe, T. E. Cheatham, III, PTRAJ and CPPTRAJ: Software for processing and analysis of molecular dynamics trajectory data. J Chem Theory Comput 9, 3084–3095 (2013).

52. Stanford University. Available at: https://getcontacts.github.io/.

53. Schrödinger L, DeLano W, PyMOL. (2020). Available from: http://www.pymol.org/pymol

