## Supplemental figures and tables for "Structural basis for conserved and distinct antigen recognition by a lineage of malaria-protective antibodies"

**Figure S1. Buried surface area (BSA) contributions of Fab 7160 CDRs to different PfCSP regions.**

The bar plot shows the relative BSA contributed by Fab 7160 CDRs H2, H3, L1, and L3 in complexes with the junctional region (magenta), minor repeat region (blue), and major repeat region (salmon). The buried surface area (BSA) contributed by each CDR to the Fab7160-peptide interaction is shown in the table.

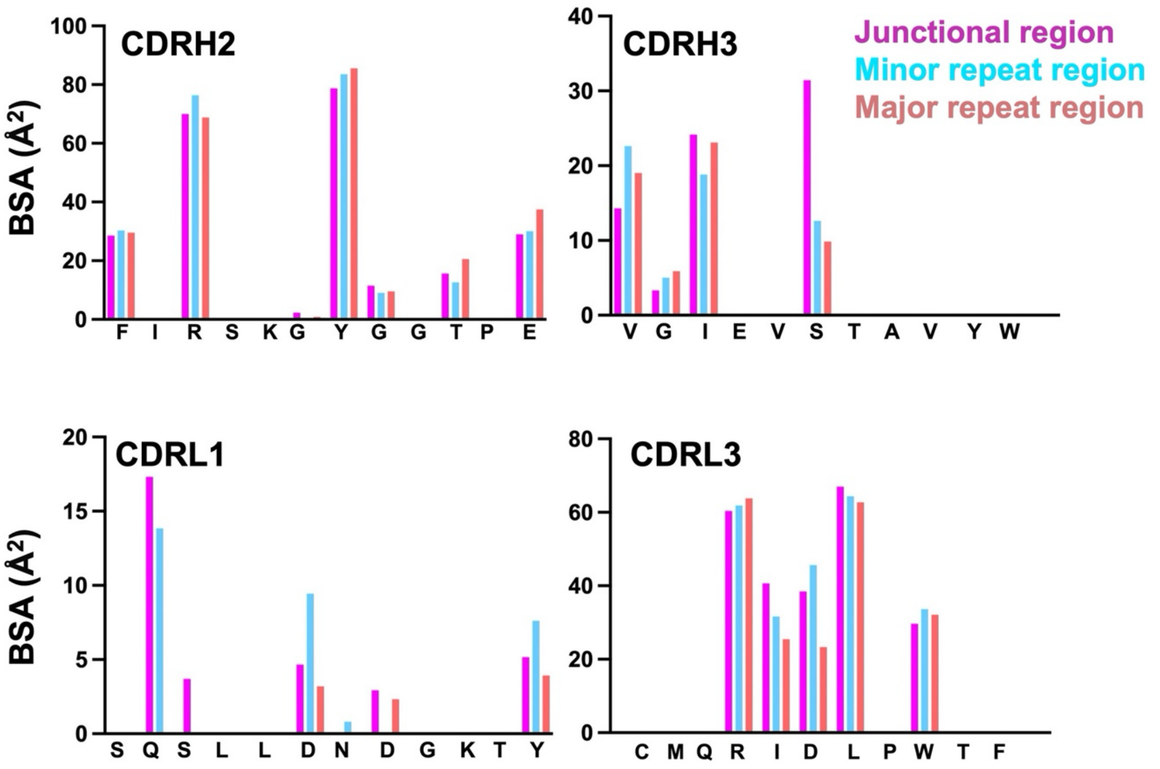

**BSA in Fab 7160-peptide complex contributed by each CDR**

| Peptide region | CDRH2 | CDRH3 | CDRL1 | CDRL3 |
|---|---|---|---|---|
| Junctional | 235 Å$^2$ | 72 Å$^2$ | 34 Å$^2$ | 234 Å$^2$ |
| Minor repeat | 232 Å$^2$ | 58 Å$^2$ | 32 Å$^2$ | 235 Å$^2$ |
| Short major repeat | 252 Å$^2$ | 58 Å$^2$ | 10 Å$^2$ | 205 Å$^2$ |

**Figure S2**. **Structural analysis of CSP peptides bound to 7160 Fab**.

The structures of the junctional (magenta), minor repeat (cyan), short major repeat (salmon), and long major repeat (yellow) peptides in the complexes with 7160 Fab are represented as sticks. Type I β-turns (black circle) and Asn pseudo $3_{10}$ turns (red circle) in the bound peptides are highlighted. Hydrogen bonds are shown as black dashes. The lower panel shows the Ramachandran plots (1) for the dihedral angles of the various peptides bound to 7160 Fab.

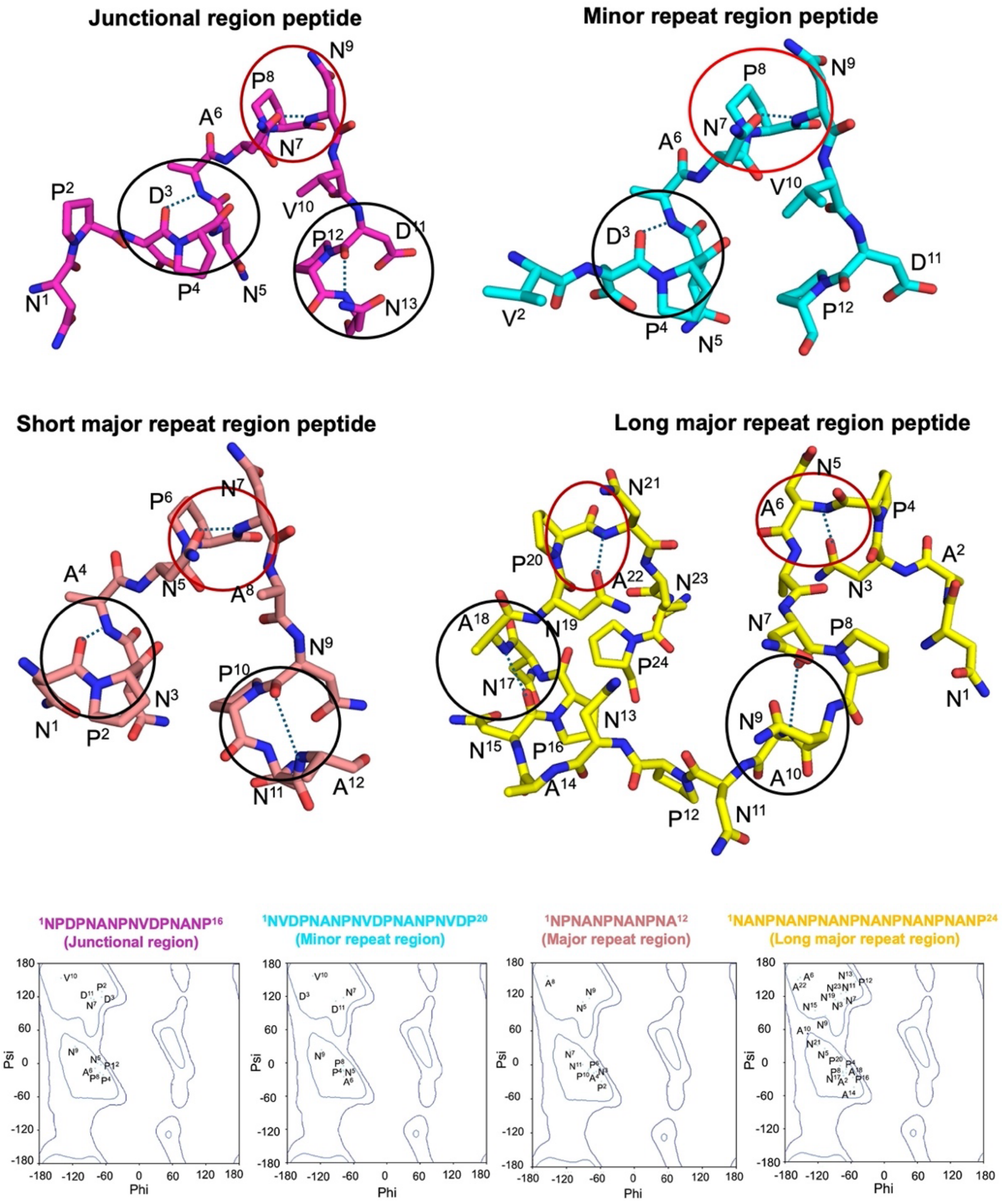

**Figure S3. Hydrogen-bond interactions of Fab 7160 and 399 with CSP-derived $NANP_6$ peptide and analysis of Fab residues in the homotypic Fab-Fab interface**

**(A-B)** Hydrogen-bond interactions between Fab 7160 (grey and pink) and Fab 399 (dark blue and pink) with $NANP_6$ (yellow). Contacts between the two Fab molecules in the homotypic Fab-Fab interaction that simultaneously recognize the peptide are highlighted. **(C-D)** A bar plot displays individual residue contributions to the buried surface area (BSA) at the homotypic Fab-Fab interface for the heavy and light chains of Fab 7160 (grey) and Fab 399 (blue). CDRs are highlighted in red in Kabat designation. Additionally, sequence alignment of the heavy and light chains with the germline gene is shown to identify somatically mutated residues. Residues involved in hydrogen bonds and salt bridges are annotated with "H" and "S," respectively.

.

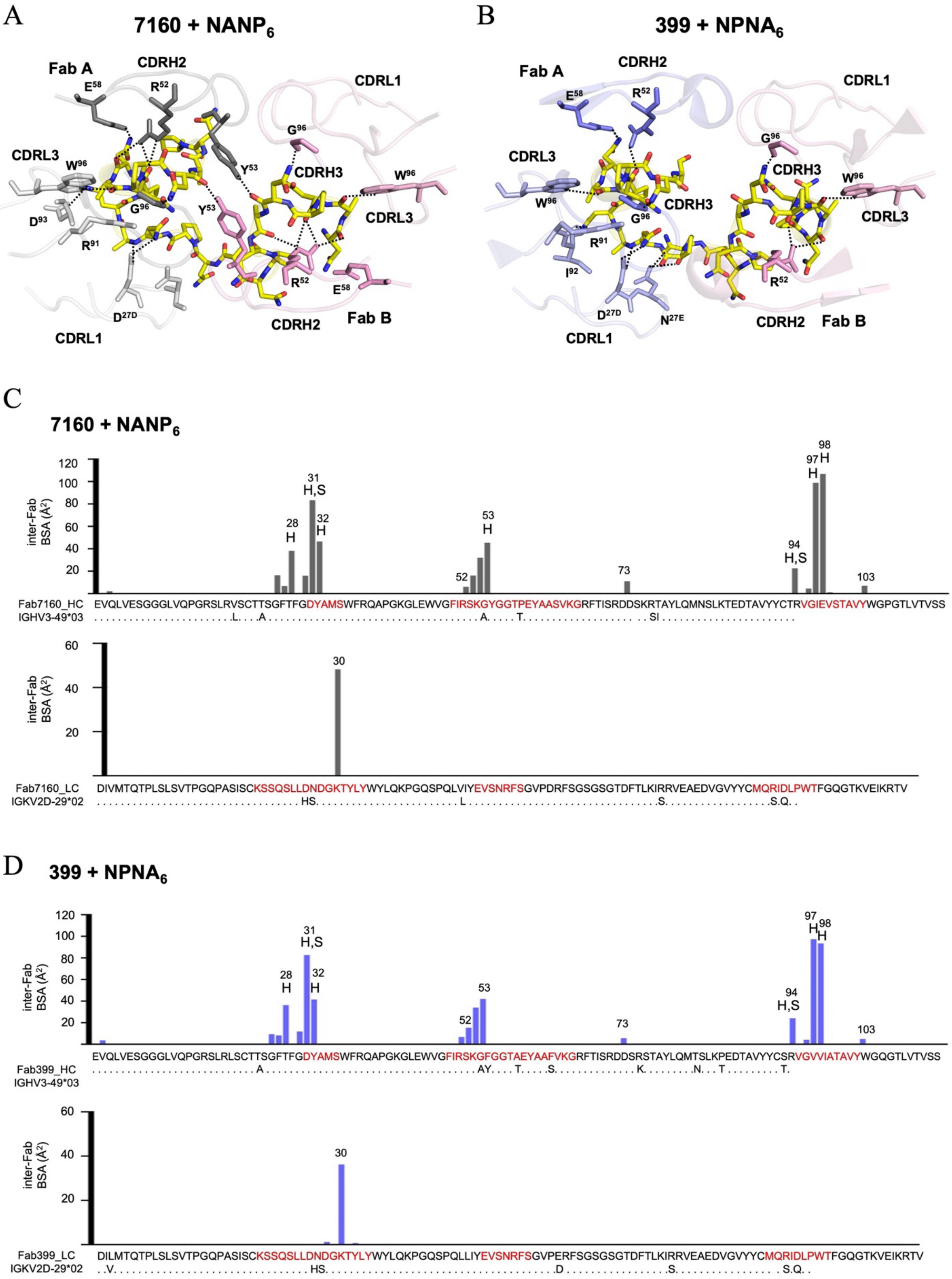
A
7160 + NANP6
Fab A
CDRH2
CDRL1
CDRL3
CDRH3
Fab B
CDRL1
CDRH2
B
399 + NPNA6
Fab A
CDRH2
CDRL1
CDRL3
CDRH3
CDRH2
Fab B
CDRL1
C
7160 + NANP6
inter-Fab BSA (Å2)
Fab7160_HC
EVQLVESGGGLVQPGRSLRVSCTTSGFTFGDYAMSWFRQAPGKGLEWVGFIRSKGYGGTPEYAASVKGRFTISRDDSKRTAYLQMNSLKTEDTAVYYCTRVGIEVSTAVYWGPGTLVTVSS
IGHV3-49*03
Fab7160_LC
DIVMTQTPLSLSVTPGQPASISCKSSQSLLDNDGKTYLYWYLQKPGQSPQLVIYEVSNRFSGVPDRFSGSGSGTDFTLKIRRVEAEDVGVYYCMQRIDLPWTFGQGTKVEIKRTV
IGKV2D-29*02
D
399 + NPNA6
inter-Fab BSA (Å2)
Fab399_HC
EVQLVESGGGLVQPGRSLRLSCTTSGFTFGDYAMSWFRQAPGKGLEWVGFIRSKGFGGTAEYAAFVKGRFTISRDDSRSTAYLQMTSLKPEDTAVYYCSRVGVVIATAVYWGQGTLVTVSS
IGHV3-49*03
Fab399_LC
DILMTQTPLSLSVTPGQPASISCKSSQSLLDNDGKTYLYWYLQKPGQSPQLLIYEVSNRFSGVPERFSGSGSGTDFTLKIRRVEAEDVGVYYCMQRIDLPWTFGQGTKVEIKRTV
IGKV2D-29*02

**Figure S4. Cryo-EM structure of Fab 7160-rsCSP**

**(A)** Side-by-side comparison of the x-ray structure of Fab 7160 bound to $NANP_6$ with the corresponding region of the cryo-EM structure of the 7160–rsCSP complex. The CSP-derived $NANP_6$ peptide is shown in yellow, and rsCSP is shown in cyan, represented as tubes and as sticks. The N and C-termini of the peptides shown are indicated. **(B)** Representative 2D class of higher-order oligomers averages observed by cryo-EM single-particle analysis. The averaged particle images, grouped during data processing, illustrate a population of two Fab 7160-dimers binding to the same rsCSP and resulting in dimer-of-dimers complexes. The head-to-head interfaces are observed in close proximity to one another in the center of the 2D classes, with the $C_H1$ constant domains for each Fab pointing outwards away from the complexes. The lack of alternative views precluded the generation of 3D models from these particles.

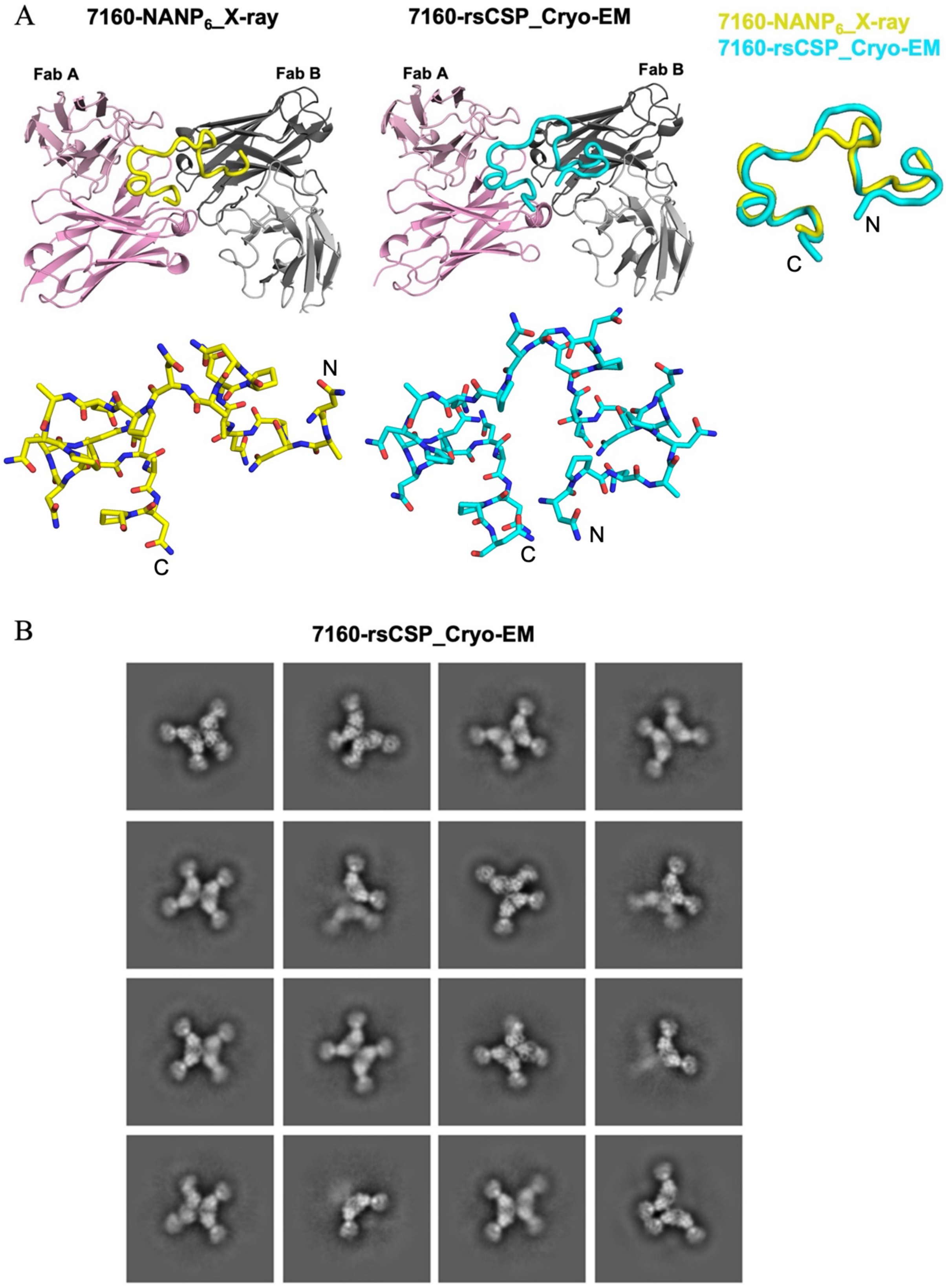
A
7160-NANP6_X-ray
7160-rsCSP_Cryo-EM
7160-NANP6_X-ray
7160-rsCSP_Cryo-EM
Fab A
Fab B
Fab A
Fab B
N
C
N
C
C
N
B
7160-rsCSP_Cryo-EM

**Figure S5. Structural comparison of Fabs 399, 7160, and 7118 highlights distinct recognition of NANP peptides**

**(A)** CDRH3 regions of Fab 399 (blue), 7160 (grey), and Fab 7118 (salmon) are shown in stick representation. **(B-C)** Superposition of CDRH3 regions of Fab 399, 7160, and 7118 indicates that the elongated CDRH3 of Fab 7118 may interfere with Fab-Fab homotypic interactions. **(D)** Structural alignment of Fab–$NANP_6$ complexes reveals distinct peptide binding modes for Fab 7160 (yellow) and Fab 7118 (magenta). The N- and C-termini of the bound peptides are indicated.

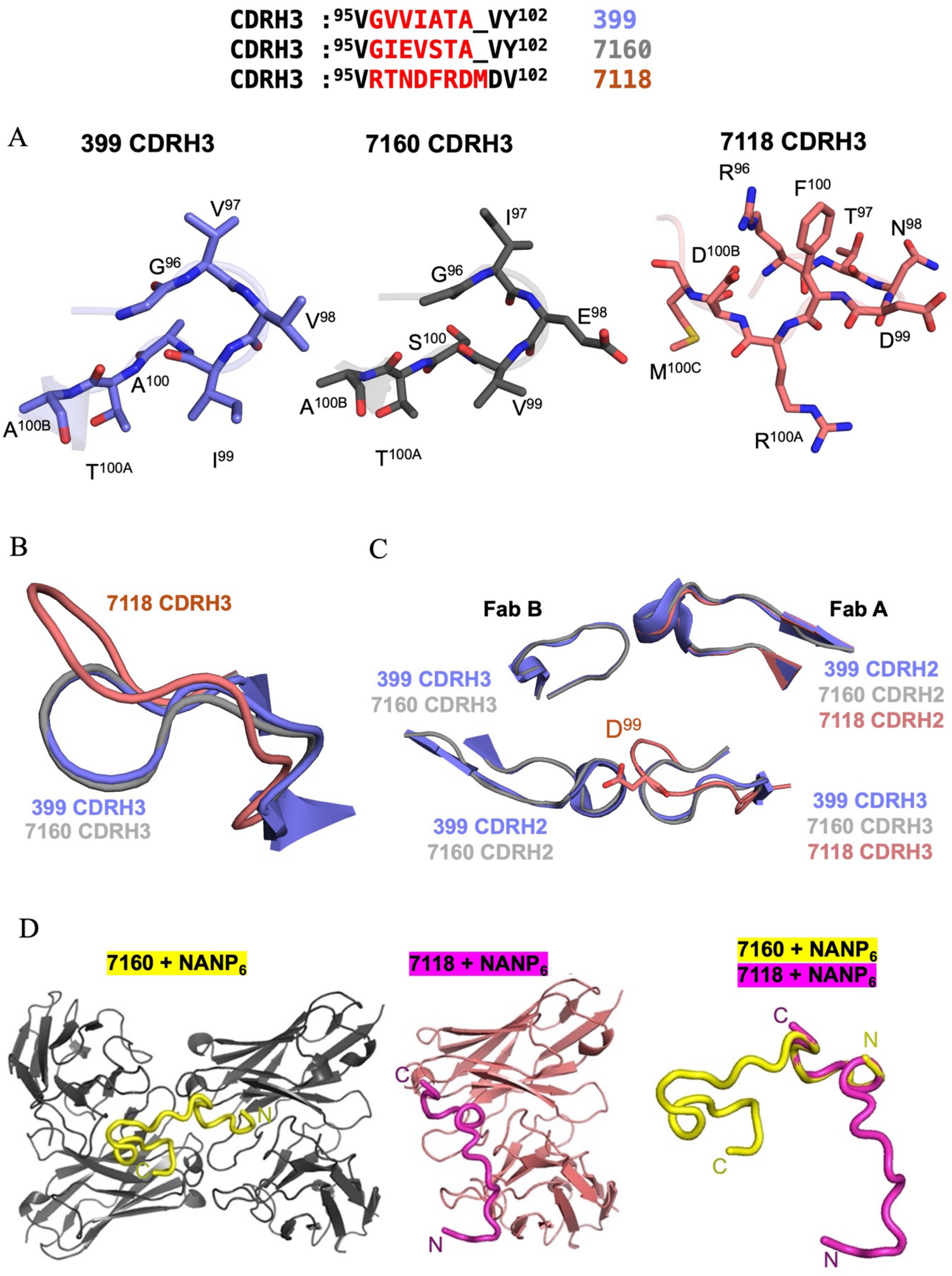

CDRH3 :95VGVVIATA_VY102 399
CDRH3 :95VGIEVSTA_VY102 7160
CDRH3 :95VRTNDFRDMDV102 7118
A
399 CDRH3
7160 CDRH3
7118 CDRH3
V97
G96
V98
A100
A100B
T100A
I99
I97
G96
E98
S100
A100B
V99
T100A
R96
F100
T97
N98
D100B
D99
M100C
R100A
B
7118 CDRH3
399 CDRH3
7160 CDRH3
C
Fab B
Fab A
399 CDRH3
7160 CDRH3
399 CDRH2
7160 CDRH2
7118 CDRH2
D99
399 CDRH3
7160 CDRH3
7118 CDRH3
399 CDRH2
7160 CDRH2
D
7160 + NANP6
7118 + NANP6
7160 + NANP6
7118 + NANP6
N
C
C
N
C
N
C
N

**Figure S6. Structural comparison of Fab 7118 with Fab 7160 and Fab 399 binding**

Superposition of Fab variable regions from two molecules of 7118-rsCSP cryo-EM structure (Fab A: green, Fab B: pink) with two molecules of Fab 7160-$NANP_6$ crystal structure (grey) in the upper panel and two molecules of Fab 399-$NPNA_6$ crystal structure (blue) shown in the lower panel. Models were aligned using PyMOL. Cartoon representation of the alignment between two 7118-rsCSP Fv molecules and the 7160-$NANP_6$ complex. $Asp^{99}$ in the CDRH3 region in 7118 is highlighted and represented as a stick. This residue would appear to introduce steric hindrance at the inter-Fab interface, disfavoring the symmetric Fab–Fab interactions observed in the Fab 7160 and 399 complexes.

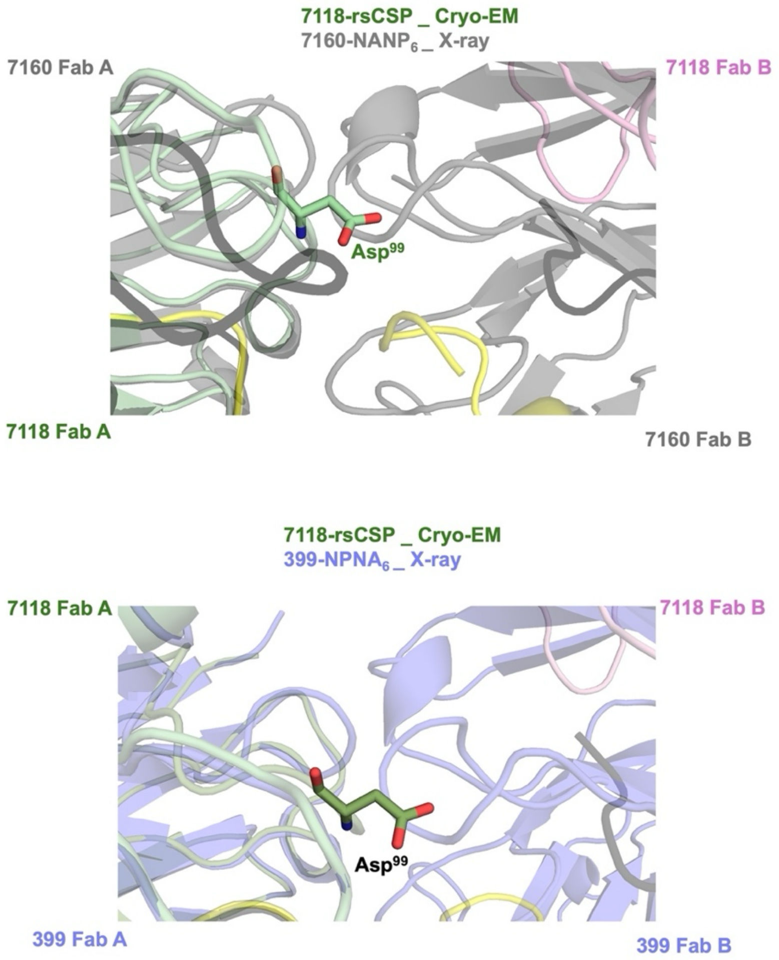

### Table S1. Biolayer interferometry kinetics of Fab binding to CSP-derived peptides

| Fab | $K_D$ (M) | $K_D$ Error | $k_{on}$ (1/Ms) | $k_{on}$ Error | $k_{off}$ (1/s) | $k_{off}$ Error |
|---|---|---|---|---|---|---|
| **NPDPNANPNVDPNANP (Junctional region)** | | | | | | |
| 399 | 2.04E-07 | 5.65E-10 | 7.19E+03 | 1.45E+01 | 1.47E-03 | 2.77E-06 |
| 7160 | 7.01E-07 | 1.45E-09 | 8.55E+03 | 1.66E+01 | 5.99E-03 | 4.25E-06 |
| 7118 | 6.12E-07 | 3.21E-09 | 2.86E+04 | 1.46E+02 | 1.75E-02 | 1.97E-05 |
| **NVDPNANPNVDPNANPNVDP (Minor repeat region)** | | | | | | |
| 399 | 2.42E-07 | 6.07E-10 | 1.04E+04 | 2.27E+01 | 2.53E-03 | 3.17E-06 |
| 7160 | 7.48E-07 | 1.75E-09 | 1.22E+04 | 2.72E+01 | 9.16E-03 | 6.75E-06 |
| 7118 | 4.84E-08 | 1.14E-10 | 3.15E+04 | 5.82E+01 | 1.53E-03 | 2.27E-06 |
| **$NPNA_3$ (Short major repeat region)** | | | | | | |
| 399 | 2.57E-08 | 7.64E-11 | 3.29E+04 | 5.29 E+01 | 8.85E-04 | 2.12E-06 |
| 7160 | 4.16E-08 | 1.02E-10 | 7.34E+04 | 1.65E+02 | 3.05E-03 | 3.06E-06 |
| 7118 | 1.07E-08 | 3.87E-11 | 4.55E+04 | 6.52E+01 | 4.89E-04 | 1.62E-06 |
| **$NANP_6$ (Long major repeat region)** | | | | | | |
| 399 | 1.06E-08 | 3.98E-11 | 8.61E+04 | 1.94E+01 | 9.16E-04 | 2.74E-06 |
| 7160 | 8.03E-09 | 2.20E-11 | 8.26E+04 | 1.14E+02 | 6.64E-04 | 1.60E-06 |
| 7118 | 5.71E-09 | 1.51E-11 | 8.89E+04 | 9.57E+01 | 5.08E-04 | 1.23E-06 |

### Table S2. X-ray data collection and refinement statistics for 7160 Fab with CSP-derived peptides

| | 7160 + junctional region | 7160 + Minor repeat region | 7160 + Short major repeat region | 7160 + Long major repeat region |
|---|---|---|---|---|
| **Data Collection** | | | | |
| Beamline | ALS 5.0.2 | ALS 5.0.2 | NSLS-II AMX | NSLS-II AMX |
| Wavelength (Å) | 0.97946 | 0.97946 | 0.92010 | 0.92010 |
| Resolution (Å) | 30.00-2.30 (2.34-2.30)[a] | 50.00-2.27 (2.31-2.27)[a] | 50.00-1.90 (1.93-1.90)[a] | 50.00-2.33 (2.39-2.33)[a] |
| Space group | $P2_12_12_1$ | $P2_12_12_1$ | $C2_1$ | $P2_1$ |
| Unit cell a, b, c (Å) | 88.19, 105.13, 405.18 | 87.00, 101.04, 104.84 | 188.01, 52.35, 102.98 | 103.21, 73.13, 134.37 |
| α, β, γ (°) | 90, 90, 90 | 90, 90, 90 | 90, 110.1, 90 | 90, 102.2, 90 |
| Unique reflections | 168,500 (8,359)[a] | 42,381 (2,085)[a] | 75,561 (3,573)[a] | 82,863 (4,058)[a] |
| Redundancy | 12.0 (11.3)[a] | 6.3 (6.1)[a] | 6.2 (4.3)[a] | 6.6 (6.3)[a] |
| Completeness (%) | 99.9 (99.9)[a] | 97.7 (98.0)[a] | 99.7 (98.1)[a] | 99.9 (99.1)[a] |
| Mean I/sigma ($\sigma_I$) | 15.1 (1.7)[a] | 14.7 (2.2)[a] | 11.4 (1.3)[a] | 13.1 (2.8)[a] |
| $R_{sym}$ (%)[b] | 18.8 (103.9)[a] | 16.0 (59.3)[a] | 14.7 (54.7)[a] | 14.9 (44.8)[a] |
| $R_{pim}$ (%)[b] | 5.6 (32.1)[a] | 6.8 (25.7)[a] | 6.4 (27.4)[a] | 6.2 (18.6)[a] |
| $CC_{1/2}$ (%)[c] | 98.9 (73.5)[a] | 99.5 (89.1)[a] | 98.5 (68.8)[a] | 98.3 (88.6)[a] |
| **Refinement statistics** | | | | |
| Resolution (Å) | 29.88-2.30 | 46.53-2.27 | 48.37-1.90 | 41.74-2.33 |
| Reflections (work) | 167,046 | 42,148 | 75,043 | 81,800 |
| Reflections (test) | 2,000 | 2,007 | 1,999 | 1,999 |
| $R_{cryst}$[d] / $R_{free}$[e] (%) | 22.8/27.4 | 25.1/28.8 | 20.7/24.5 | 20.6/23.6 |
| **Number of atoms** | | | | |
| Fab | 26,634 | 6,683 | 6,636 | 13,318 |
| Peptide | 718 | 160 | 168 | 316 |
| Water | 338 | 120 | 250 | 275 |
| **Average B-value (Å$^2$)** | | | | |
| Fab | 43 | 37 | 31 | 33 |
| Peptide | 44 | 38 | 30 | 37 |
| Water | 36 | 37 | 32 | 31 |
| Wilson B (Å$^2$) | 38 | 34 | 23 | 30 |
| **RMSD from ideal geometry** | | | | |
| Bond angle (°) | 0.55 | 0.54 | 0.93 | 0.53 |
| Bond length (Å) | 0.002 | 0.002 | 0.008 | 0.002 |
| **Ramachandran statistics**[f] | | | | |
| Favored (%) | 98.13 | 97.51 | 98.64 | 98.64 |
| Allowed (%) | 1.87 | 2.49 | 1.36 | 1.36 |
| Outliers (%) | 0.00 | 0.00 | 0.00 | 0.00 |
| **PDB Code** | 9ZM7 | 9ZM8 | 9ZM9 | 9ZMA |

[a] Numbers in parentheses refer to the highest resolution shell.
[b] Rsym = Σhkl Σi | Ihkl,i - | / Σhkl Σi Ihkl,i and Rpim = Σhkl (1/(n-1))1/2 Σi | Ihkl,i - | / Σhkl Σi Ihkl,i, where Ihkl,i is the scaled intensity of the ith measurement of reflection h, k, l, is the average intensity for that reflection, and n is the redundancy.
[c] CC1/2 = Pearson correlation coefficient between two random half datasets.
[d] Rcryst = Σhkl | Fo - Fc | / Σhkl | Fo | x 100, where Fo and Fc are the observed and calculated structure factors, respectively.
[e] Rfree was calculated as for Rcryst, but on a test set comprising 5% of the data excluded from refinement.
[f] From MolProbity (2).

### Table S3. X-ray data collection and refinement statistics for 7118 Fab with CSP-derived peptides

| | **7118 + Minor repeat region** | **7118 + Long major repeat region** |
|---|---|---|
| **Data Collection** | | |
| Beamline | SSRL 12-1 | SSRL 12-1 |
| Wavelength (Å) | 0.97946 | 0.97946 |
| Resolution (Å) | 50.00-2.09 (2.14-2.09)[a] | 50.00-1.99 (2.03-1.99)[a] |
| Space group | $P2_12_12_1$ | $P2_12_12_1$ |
| Unit cell a, b, c (Å) | 73.84, 83.03, 88.46 | 74.15, 83.39, 89.55 |
| α, β, γ (°) | 90, 90, 90 | 90, 90, 90 |
| Unique reflections | 30,491 (1,561)[a] | 37,945 (1,871)[a] |
| Redundancy | 5.1 (6.3)[a] | 6.0 (6.1)[a] |
| Completeness (%) | 93.7 (98.5)[a] | 99.5 (99.7)[a] |
| Mean I/sigma ($\sigma_I$) | 29.0 (4.7)[a] | 38.6 (7.4)[a] |
| $R_{sym}$ (%)[b] | 8.8 (47.2)[a] | 8.3 (34.0)[a] |
| $R_{pim}$ (%)[b] | 4.7 (20.3)[a] | 3.8 (15.3)[a] |
| $CC_{1/2}$ (%)[c] | 99.9 (86.8)[a] | 99.9 (94.1)[a] |
| **Refinement statistics** | | |
| Resolution (Å) | 37.94-2.09 | 41.70-2.00 |
| Reflections (work) | 30,283 | 37,842 |
| Reflections (test) | 1,999 | 2,002 |
| $R_{cryst}$[d] / $R_{free}$[e] (%) | 20.6/24.2 | 18.0/21.7 |
| **Number of atoms** | | |
| Fab | 3,337 | 3,346 |
| Peptide | 138 | 140 |
| Water | 133 | 251 |
| **Average B-value (Å²)** | | |
| Fab | 28 | 23 |
| Peptide | 31 | 23 |
| Water | 30 | 29 |
| Wilson B (Å²) | 38 | 23 |
| **RMSD from ideal geometry** | | |
| Bond angle (°) | 0.59 | 0.90 |
| Bond length (Å) | 0.002 | 0.007 |
| **Ramachandran statistics[f]** | | |
| Favored (%) | 98.43 | 98.00 |
| Allowed (%) | 1.57 | 1.78 |
| Outliers (%) | 0.00 | 0.22 |
| **PDB Code** | 9ZMB | 9ZMC |

[a] Numbers in parentheses refer to the highest resolution shell.
[b] Rsym = Σhkl Σi | Ihkl,i - | / Σhkl Σi Ihkl,i and Rpim = Σhkl (1/(n-1))1/2 Σi | Ihkl,i - | / Σhkl Σi Ihkl,i, where Ihkl,i is the scaled intensity of the ith measurement of reflection h, k, l, is the average intensity for that reflection, and n is the redundancy.
[c] CC1/2 = Pearson correlation coefficient between two random half datasets.
[d] Rcryst = Σhkl | Fo - Fc | / Σhkl | Fo | x 100, where Fo and Fc are the observed and calculated structure factors, respectively.
[e] Rfree was calculated as for Rcryst, but on a test set comprising 5% of the data excluded from refinement.
[f] From MolProbity (2).

### Table S4A. Hydrogen bonds between Fab 7160 and (NANP)$_6$ peptide with BSA.

| NANP$_6$ (BSA Å$^2$) | Distance (Å) | 7160-HC Fab A | 7160-LC Fab A | 7160-HC Fab B | 7160-LC Fab B |
|---|---|---|---|---|---|
| **Ala2 (92)** | | | | | |
| Ala-O | 3.33 | | Trp$^{96}$-NE1 | | |
| **Asn3 (34)** | | | | | |
| Asn-O | 2.90 | Arg$^{52}$-NH2 | | | |
| Asn-O | 3.10 | Arg$^{52}$-NE | | | |
| **Asn5 (124)** | | | | | |
| Asn-ND2 | 2.97 | Gly$^{96}$-O | | | |
| Asn-O | 2.83 | | | Tyr$^{53}$-OH | |
| **Ala6 (17)** | | | | | |
| Ala-O | 3.06 | Arg$^{52}$-NE | | | |
| **Pro12 (11)** | | | | | |
| Pro-O | 3.60 | | | | Asn$^{27E}$-ND2 |
| **Ala14 (49)** | | | | | |
| Ala-N | 3.16 | | | | Asp$^{27D}$-OD1 |
| **Asn15 (47)** | | | | | |
| Asn-ND2 | 2.85 | | | | Asp$^{93}$-OD1 |
| **Asn-17 (45)** | | | | | |
| Asn-ND2 | 2.94 | | | Glu$^{58}$-OE2 | |
| Asn-O | 2.92 | | | Arg$^{52}$-NH2 | |
| **Ala18 (68)** | | | | | |
| Ala18-O | 3.38 | | | | Arg$^{91}$-NE |
| Ala18-O | 3.24 | | | | Trp$^{96}$-NE1 |
| **Asn19 (21)** | | | | | |
| Asn-OD1 | 3.32 | | | | Arg$^{91}$-NH1 |
| Asn-O | 2.99 | | | Arg$^{52}$-NE | |
| Asn-O | 3.21 | | | Arg$^{52}$-NH2 | |
| **Asn21 (123)** | | | | | |
| Asn-O | 3.19 | Tyr$^{53}$-OH | | | |
| Asn-ND2 | 2.96 | | | Gly$^{96}$-O | |

### Table S4B. Hydrogen bonds between Fab 7118 and (NANP)$_6$ peptide with BSA

| NANP$_6$ (BSA Å$^2$) | Distance (Å) | 7118-HC | 7118-LC |
|---|---|---|---|
| **Asn5 (50)** | | | |
| Asn-O | 2.86 | | Ser$^{27A}$-OG |
| **Asn7 (102)** | | | |
| Asn-N | 2.76 | | Leu$^{27C}$-O |
| Asn-ND2 | 3.33 | | Glu$^{27}$-OE1 |
| Asn-ND2 | 2.92 | | Ser$^{27A}$-O |
| **Pro8 (21)** | | | |
| Pro-O | 2.75 | | Arg$^{27E}$-NH2 |
| **Ala10 (51)** | | | |
| Ala-O | 2.77 | | His$^{27D}$-ND2 |
| **Asn11 (50)** | | | |
| Asn-ND2 | 3.24 | | Ile$^{92}$-O |
| Asn-ND2 | 3.40 | | Asp$^{93}$-OD1 |
| **Asn13 (48)** | | | |
| Asn-O | 2.99 | Arg$^{52}$-NH2 | |
| Asn-ND2 | 2.66 | Glu$^{58}$-OE2 | |
| **Ala14 (60)** | | | |
| Ala-O | 3.25 | | Trp$^{96}$-NE1 |
| **Asn15 (33)** | | | |
| Asn-O | 2.87 | Arg$^{52}$-NE | |
| **Asn17 (118)** | | | |
| Asn-O | 3.30 | Asn$^{53}$-ND2 | |
| Asn-ND2 | 2.93 | Arg$^{96}$-O | |
| Asn-ND2 | 3.07 | Asn$^{98}$-O | |
| Asn-ND2 | 2.88 | Phe$^{100}$-O | |

### Table S5. Cryo-EM structure data collection and refinement statistics of Fab 7118-rsCSP

| | **7118 -rsCSP EMDB-74167** | **7160-rsCSP EMDB-74165** |
|---|---|---|
| **Data collection and processing** | | |
| Magnification | 190,000x | 190,000x |
| Voltage (kV) | 200 | 200 |
| Electron exposure ($e^{-}/Å^2$) | 60 | 60 |
| Defocus range (μm) | -0.6 to -1.6 | -0.6 to -1.6 |
| Pixel size (Å) | 0.718 | 0.718 |
| Symmetry imposed | C1 | C1 |
| Final particle images (no.) | 250,774 | 116,764 |
| Map resolution (Å) | 3.32 | 3.55 |
| FSC threshold | 0.143 | 0.143 |
| **Refinement statistics** | | |
| Map sharpening *B* factor ($Å^2$) | -108 | -93 |
| **Model composition** | | |
| Non-hydrogen atoms | 7576 | 6857 |
| Protein residues | 981 | 902 |
| ***Average B* values ($Å^2$)** | | |
| Fab | 69 | 78 |
| CSP | 57 | 67 |
| **RMSD** | | |
| Bond angle (°) | 0.65 | 0.629 |
| Bond length (Å) | 0.005 | 0.004 |
| **Validation** | | |
| MolProbity score | 1.82 | 1.18 |
| Clashscore | 6.37 | 2.95 |
| Poor rotamers (%) | 0.00 | 0.00 |
| **Ramachandran statistics** | | |
| Favored (%) | 92.52 | 97.52 |
| Allowed (%) | 7.48 | 2.48 |
| Disallowed (%) | 0.00 | 0.00 |
| **PDB Code** | 9ZFZ | 9ZFY |

**Table S6. Mean electrostatic energy of 7118 Fab-rsCSP interfaces calculated during molecular dynamic simulations.**

| **Interface** | **ELE (kcal/mol, mean ± std)** |
|---|---|
| Fab A - CSP | -120.37 ± 27.41 |
| Fab B - CSP | -122.10 ± 45.08 |
| Fab C - CSP | -139.64 ± 43.97 |
| Fab D – CSP | -59.82 ± 24.98 |
| All Fabs - CSP | -441.92 ± 68.84 |

### References

1. M. Kumar, R. S. Rathore, RamPlot: a webserver to draw 2D, 3D and assorted Ramachandran (φ,ψ ) maps. *J Appl Crystallogr.* **58**, 630-636 (2025).

2. V. B. Chen *et al.*, MolProbity: all-atom structure validation for macromolecular crystallography. *Acta Crystallogr D Biol Crystallogr* **66**, 12-21 (2010).
